# CellPhy: accurate and fast probabilistic inference of single-cell phylogenies from scDNA-seq data

**DOI:** 10.1101/2020.07.31.230292

**Authors:** Alexey Kozlov, Joao M. Alves, Alexandros Stamatakis, David Posada

## Abstract

We introduce a maximum likelihood framework called CellPhy for inferring phylogenetic trees from single-cell DNA sequencing (scDNA-seq) data. CellPhy leverages a finite-site Markov genotype substitution model with 16 diploid states, akin to those typically used in statistical phylogenetics. It includes a dedicated error function for single cells that incorporates amplification/sequencing error and allelic dropout (ADO). Moreover, it can explicitly consider the uncertainty of the variant calling process by using genotype likelihoods as input. We implemented CellPhy in a widely used open-source phylogenetic inference package (RAxML-NG) that provides statistical confidence measurements on the estimated tree and scales particularly well on large scDNA-seq datasets with hundreds or thousands of cells. To benchmark CellPhy, we carried out 19,400 coalescent simulations of cell samples from exponentially-growing tumors for which the true phylogeny was known. We evolved single-cell diploid DNA genotypes along the simulated genealogies under different scenarios, including infinite- and finite-sites nucleotide mutation models, trinucleotide mutational signatures, sequencing, and amplification errors, allele dropouts, and cell doublets. Our simulations suggest that CellPhy is robust to amplification/sequencing errors and ADO and outperforms state-of-the-art methods under realistic scDNA-seq scenarios both in terms of accuracy and speed. Also, we sequenced 24 single-cell whole-genomes from a colorectal tumor. Together with three published scDNA-seq data sets, we analyzed these empirical data to illustrate how CellPhy can provide more reliable biological insights than most competing methods. CellPhy is freely available at https://github.com/amkozlov/cellphy.

## Introduction

The study of single cells is revolutionizing biology, unveiling unprecedented levels of genomic and phenotypic heterogeneity within otherwise seemingly homogeneous tissues (Macaulay and Voet 2014; Stegle et al. 2015; Gawad et al. 2016; Baslan and Hicks 2017; Tanay and Regev 2017; Lim et al. 2020). Understanding this somatic mosaicism has applications in multiple areas of Biology due to its intrinsic connection to development, aging, and disease (Marioni and Arendt 2017; Wiedmeier et al. 2019). However, the analysis of single-cell genomic data is not devoid of challenges (Stegle et al. 2015; Lähnemann et al. 2020), including the development of more integrative, scalable, and biologically realistic models of somatic evolution that can handle the inherent noise of single-cell data –mainly amplification error and allelic dropout (ADO) (Navin 2014). Understanding how somatic cells evolve is one of the main applications of single-cell technologies. In particular, the reconstruction of cell phylogenies from single-cell DNA sequencing (scDNA-seq) can help us understand the mode and tempo of cell diversification and the mechanisms behind it. When geographical information is also available, the cell genealogy can inform us about cell expansions and migrations, which are of great relevance, for example, to understand tumor growth and metastases.

Several methods have been proposed to reconstruct phylogenetic trees from scDNA-seq data (Kuipers, Jahn, and Beerenwinkel 2017; Zafar et al. 2018). OncoNem (Ross and Markowetz 2016) implements a nested-effects likelihood model to correct for observational noise, plus a simple heuristic search that attempts to maximize the likelihood of the data across tree space. It reconstructs a tree of clones and mutations by assigning cells to the clones. OncoNem assumes an infinite-site mutation (ISM) model, and as we will see below, it is very slow and can only analyze small datasets. SCITE (Jahn et al. 2016) implements an ISM similar to the one used by OncoNem but uses Markov Chain Monte Carlo (MCMC) to sample likelihoods/posterior probabilities. It essentially estimates a mutation tree to which it attaches the cells a posteriori. Conveniently, it can infer false-positive (FP) and false-negative (FN) error rates from the data. infSCITE (Kuipers, Jahn, Raphael, et al. 2017) extends SCITE to consider cell doublets, test the ISM’s validity, and learn the FP rate from panel sequencing data. SiFit (Zafar et al. 2017) implements a Markov finite-site model of evolution and a heuristic ML tree search algorithm. It can also estimate FP and FN error rates, and in the reported simulations, outperforms OncoNem and SCITE in terms of speed and accuracy. Finally, SCIPhI (Singer et al. 2018) jointly estimates the cell genotypes and their phylogenetic relationships, working directly with the read counts and considering amplification/sequencing errors, ADO, and loss of heterozygosity.

All methods above were implemented *de novo* for the specific problem of single-cell phylogeny reconstruction. Statistical phylogenetics is a well-developed field, with many sophisticated evolutionary models implemented into efficient computational programs. We reasoned that we could leverage such a framework to obtain more accurate somatic cell phylogenies. Thus, we implemented a novel, dedicated model for single-cell somatic evolution in an existing, successful framework for statistical phylogenetics, RAxML-NG (Kozlov et al. 2019). We named the resulting software CellPhy. Our computer simulations and the analysis of several empirical datasets show that CellPhy can reconstruct more accurate single-cell phylogenies across different biological scenarios and is substantially faster than the competing likelihood-based methods. We also evaluated a maximum parsimony-based tool TNT (Goloboff and Catalano 2016), an extremely fast approach for phylogenetic reconstruction. Although TNT performed very well in error-free simulation scenarios, its accuracy quickly degraded in the presence of the typical biases observed in scDNA-seq data.

## Results

### CellPhy

We developed a probabilistic model for the phylogenetic analysis of single-cell diploid genotypes inferred from scDNA-seq experiments, called CellPhy. For tractability, we focused on single nucleotide variants (SNVs), which is arguably the most common type of genetic data obtained from somatic tissues nowadays. Current models of evolution for single-cells only consider the absence/presence of mutations regardless of the nucleotides involved (Jahn et al. 2016; Ross and Markowetz 2016; Zafar et al. 2017). Thus, they deal with at most a ternary state space where genotypes can only have 0, 1, or 2 mutant alleles (normal, heterozygous, or homozygous mutant, respectively). Instead, here we consider changes among all possible 16 phased DNA genotypes Γ = {A|A, A|C, A|G, A|T, C|A, C|C, C|G, C|T, G|A, G|C, G|G, G|T, T|A, T|C, T|G, T|T}, by extending the well-established finite-site continuous-time general-time-reversible Markov model of DNA sequence evolution with four states (GTR) (Tavaré 1986) to 16 states. We named this model GT16.

Because somatic evolution proceeds mainly by mitosis, where both daughter cells receive the same set of chromosomes and recombination can be safely ignored, there is a unique cell history recorded in the same way in the maternal and paternal chromosomes. Therefore, we do not need to know whether a mutation occurred in the maternal or paternal chromosome to infer the cell phylogeny. While the GT16 model in CellPhy considers phased genotypes, nowadays, the vast majority of the empirical scDNA-seq datasets are unphased due to technical limitations. Conveniently, the GT16 model can also work with unphased genotypes (10 states) simply by considering the ambiguity of the phase in heterozygotes (see Methods). Our simulations consist of unphased genotypes to represent current scDNA-seq data.

Importantly, single-cell genotypes can be very noisy, mainly due to biases during whole-genome amplification (WGA) (Navin 2014). While previous methods rely on observational error models based on FP and FN rate parameters, we built an error model with two free parameters, the ADO rate (*δ*) and the amplification/sequencing error rate (*ε*). Compared with previous implementations, the advantage of this parameterization is that it can incorporate plausible situations such as an amplification/sequencing error converting a homozygous mutant into a heterozygous genotype. Due to its low probability of occurrence, we discard the possibility of observing more than one amplification/sequencing error at a given site. Still, we allow for the presence of both ADO and amplification/sequencing error in a single genotype. Instead of using a genotype error model, CellPhy can also use the Phred-scaled genotype likelihoods provided by single-cell variant callers.

We assume that the evolutionary history of a sample of cells can be appropriately portrayed as an unrooted binary tree and that all SNVs evolve in the same way and independently of each other. Given a set of single-cell SNV genotypes (provided by the user as a matrix in FASTA or PHYLIP format, or as a standard VCF file), CellPhy leverages its error and genotype models to compute the tree likelihood as a product of the independent probabilities across SNVs, using the standard Felsenstein pruning algorithm (Felsenstein 1981; Felsenstein 2004). Conveniently, CellPhy does not need to assume any particular genotypic configuration at the root, like other programs. For example, SiFit assumes that the root of the tree is homozygous for the reference allele at all sites. Instead, the CellPhy tree can be easily rooted *a posteriori* using a particular set of cells as an outgroup (see Felsenstein 2004). For example, if we study tumor cells, the outgroup could be one or more healthy cells.

We implemented CellPhy’s phylogenetic model in RAxML-NG (Kozlov et al. 2019), a popular maximum likelihood (ML) framework in organismal phylogenetics. Therefore, to obtain ML estimates of the model parameters (substitution rates, ADO, and amplification/sequencing errors) and the cell tree, CellPhy leverages the optimization routines and tree search strategies of RAxML (Stamatakis 2014) and RAxML-NG (Kozlov et al. 2019). The latter, for example, is known to work particularly well on large trees (Zhou et al. 2018). The fact that CellPhy exploits RAxML-NG allows it to also calculate confidence values for the inner tree branches using either the standard (Felsenstein 1985) or transfer (Lemoine et al. 2018) bootstrap (BS) techniques. Moreover, CellPhy can perform ancestral state reconstruction (Yang et al. 1995) to obtain ancestral ML genotypes and map mutations onto the branches of the ML tree. CellPhy is freely available, together with documentation, tutorials, and example data at https://github.com/amkozlov/cellphy.

### Validation and benchmarking

#### Simulation 1: infinite-site model and low number of SNVs (“target-ISM”)

For simulated datasets of 40 cells and 250-1000 SNVs under an infinite-site mutation model, phylogenetic accuracy decreased rapidly for all methods with increasing levels of genotype error or ADO (Figures 1, S1-S2). CellPhy was the most accurate method overall, although closely followed by SiFit and, to a lesser extent, by infSCITE, which performed worse when genotype errors were prevalent. The parsimony-based inference tool TNT was as accurate as CellPhy, SiFit, or infSCITE without ADO and genotype error, but worse otherwise. OncoNEM, which produced highly unresolved trees (i.e., 50% to 90% polytomies), performed poorly under all scenarios. Because OncoNEM is also extremely slow and cannot handle large data sets (see the Computational speed section below), we did not evaluate it further. The genotype coding strategy (“keep”, “remove”, and “missing”) influenced accuracy only when genotype errors were present, with “missing” and “keep” being better than “remove” (see Figures S1-S2).

**Figure 1.**
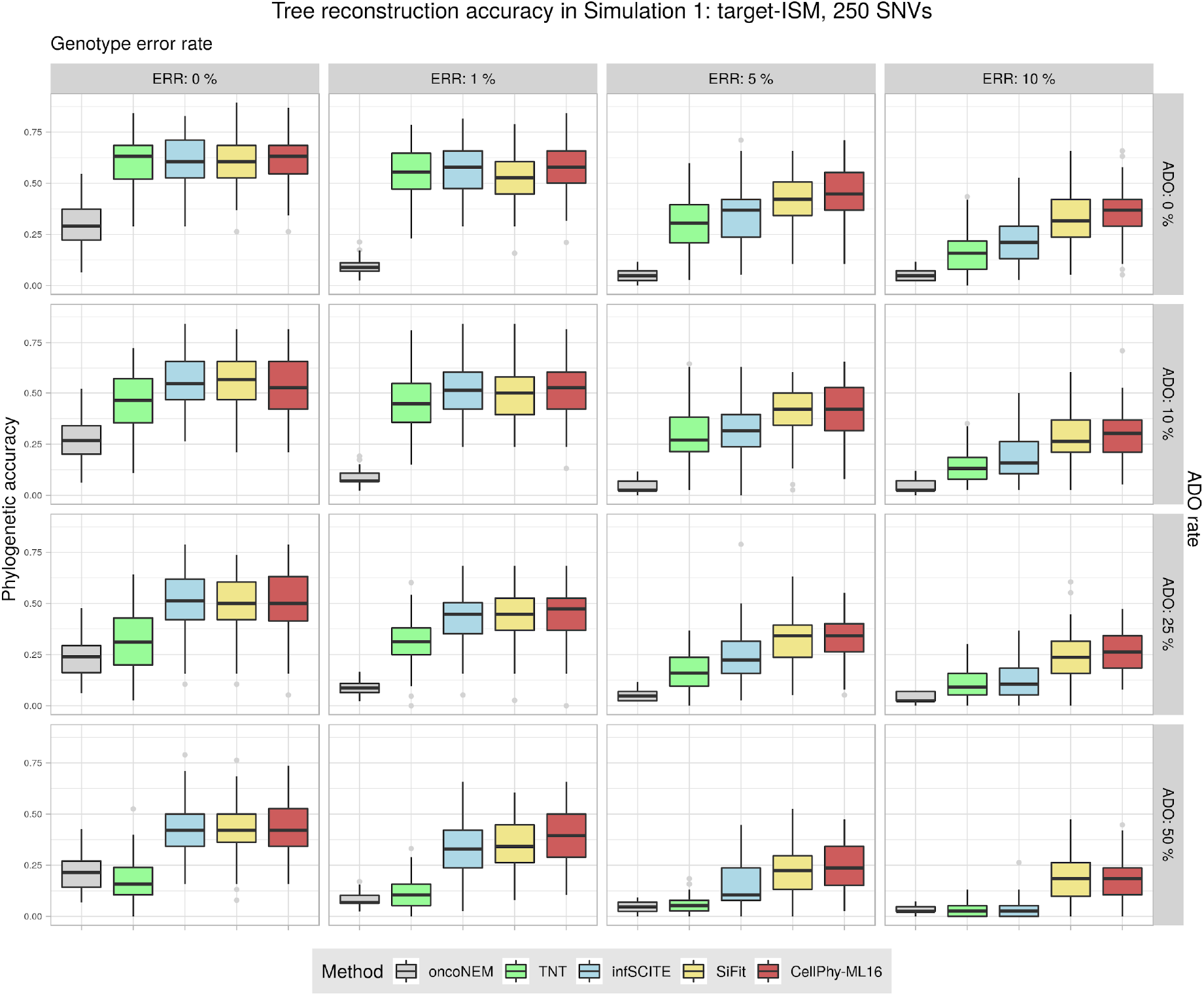
Phylogenetic accuracy in Simulation 1 (“target-ISM”) with 250 SNVs. Datasets consisted of 40 cells. Accuracy was evaluated under different levels of genotype error (ERR), allele dropout (ADO), and genotype recoding strategy “missing”. Phylogenetic accuracy is 1 – nRF (see Methods). Boxplots were generated with the ggplot2 R package (Wickham 2016) (https://ggplot2.tidyverse.org) with default parameters. Lower and upper hinges correspond to the first and third quartiles. The upper whisker extends from the hinge to the largest value no further than 1.5 * IQR from the hinge (where IQR is the interquartile range or distance between the first and third quartiles). The lower whisker extends from the hinge to the smallest value at most 1.5 * IQR of the hinge. Data points beyond the end of the whiskers are called “outlier” points and plotted individually.

#### Simulation 2: finite-site model and large number of SNVs (“WGS-FSM”)

When we simulated larger data sets (100 cells, ^~^2,000 SNVs) under a finite-site model of DNA evolution, overall phylogenetic accuracy increased for all methods. As before, all strategies performed worse with higher levels of ADO or genotype error (Figure 2). CellPhy was again most accurate overall, mainly when the data contained many genotype errors and ADO events. As in Simulation 1, the “missing” coding strategy was slightly superior to “keep”, particularly with higher genotype error rates, and substantially better than “remove” (data not shown). Thus, for subsequent simulations, we only considered the “missing” strategy.

**Figure 2.**
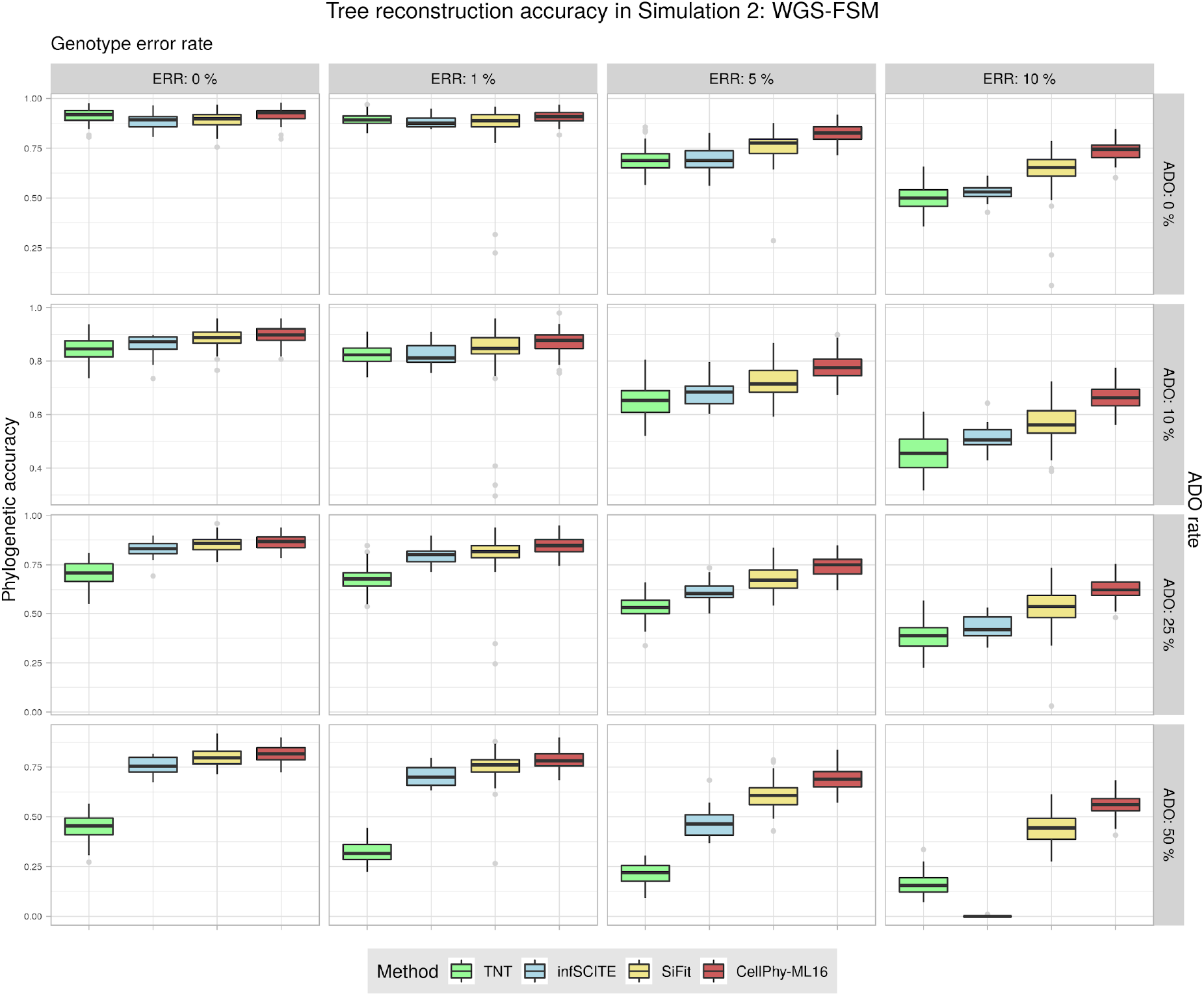
Phylogenetic reconstruction accuracy in Simulation 2 (“WGS-FSM”). Datasets consisted of 100 cells and ^~^2000 SNVs. Accuracy was evaluated under different levels of genotype error (ERR), allelic dropout (ADO), and genotype coding strategies “missing”. Phylogenetic accuracy is 1 – nRF (see Methods). See Figure 1 for an explanation of the boxplots.

#### Simulation 3: mutational signatures and large number of SNVs (“WGS-sig”)

In Simulation 3, we produced relatively large datasets (60 cells, 1,000-4,000 SNVs) under COSMIC trinucleotide mutational signatures 1 and 5 (i.e., assuming an infinite-site model). The trends were as before, and CellPhy consistently outperformed the competing methods, especially with increasing levels of genotype error or ADO (Figures S3-S4).

#### Simulation 4: genotype likelihoods from NGS read counts (“NGS-like”)

Here, we simulated NGS data, and the input for tree inference consisted of read counts (SCIPhI), ML genotypes (CellPhy-ML and the remaining methods), or genotype likelihoods (CellPhy-GL). Across this scenario, with a realistic sequencing depth for single-cells (5x), CellPhy performed much better than the competing methods, in particular under its genotype likelihood mode (“CellPhy-GL”) (Figure 3). While at 30x and 100x, the accuracy differences were substantially more minor, CellPhy still performed as well as or better than the competing methods (Figures S5-S6). Strikingly, the relative performance of SCIPhI worsens at 30x and fully degrades at 100x in the presence of errors.

**Figure 3.**
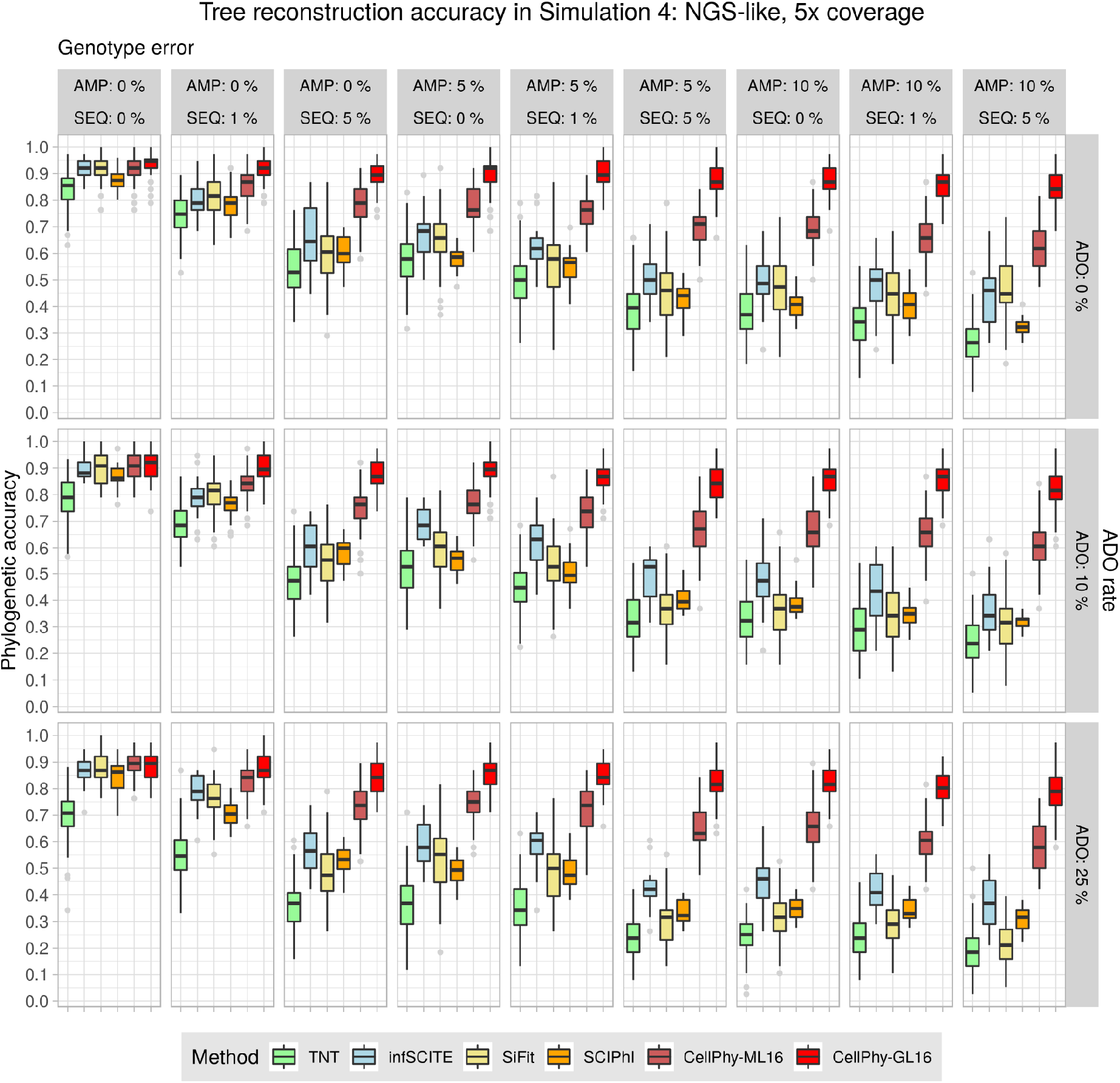
Phylogenetic accuracy in Simulation 4 (“NGS-like”) at 5x. Datasets consisted of 40 cells and 1000-2000 SNVs. TNT, SiFiT, and CellPhy-ML16 use the inferred genotypes, CellPhy-GL16 uses the genotype likelihoods, and SCIPhI uses read counts. Phylogenetic accuracy is 1 – nRF (see Methods). AMP is the amplification error rate, SEQ is the sequencing error rate, and ADO is the allelic dropout rate. CellPhy was run using the ML genotypes (CellPhy-ML) and using the genotype likelihoods (CellPhy-GL). See Figure 1 for an explanation of the boxplots.

#### Simulation 5: NGS doublets

In many single-cell experiments, depending on the isolation method used, cell doublets can be relatively common (Stegle et al. 2015), so we also assessed their effect. As expected, the presence of cell doublets reduced phylogenetic accuracy for all methods. Still, CellPhy was the most accurate method, particularly with ADO and amplification errors (Figure S7).

#### Simulation 6: NGS for large numbers of cells and SNVs

We assessed phylogenetic accuracy on large scDNA-seq datasets, with up to 1,000 cells and 50,000 SNVs, without doublets. In this case, we could not evaluate infSCITE, as jobs were still running after one month. Here, CellPhy generally outperformed the competing methods, with rapidly increasing accuracy as a function of the number of SNVs, benefiting further from the use of genotype likelihoods (Figure 4). Remarkably, SCIPhI showed constant accuracy of ^~^0.4 across conditions. We noted that the SCIPhI trees contained many unresolved nodes, which decreases the number of possible false positives. Moreover, SCIPhI’s accuracy decreased with more SNVs, highlighting a systematic bias.

**Figure 4.**
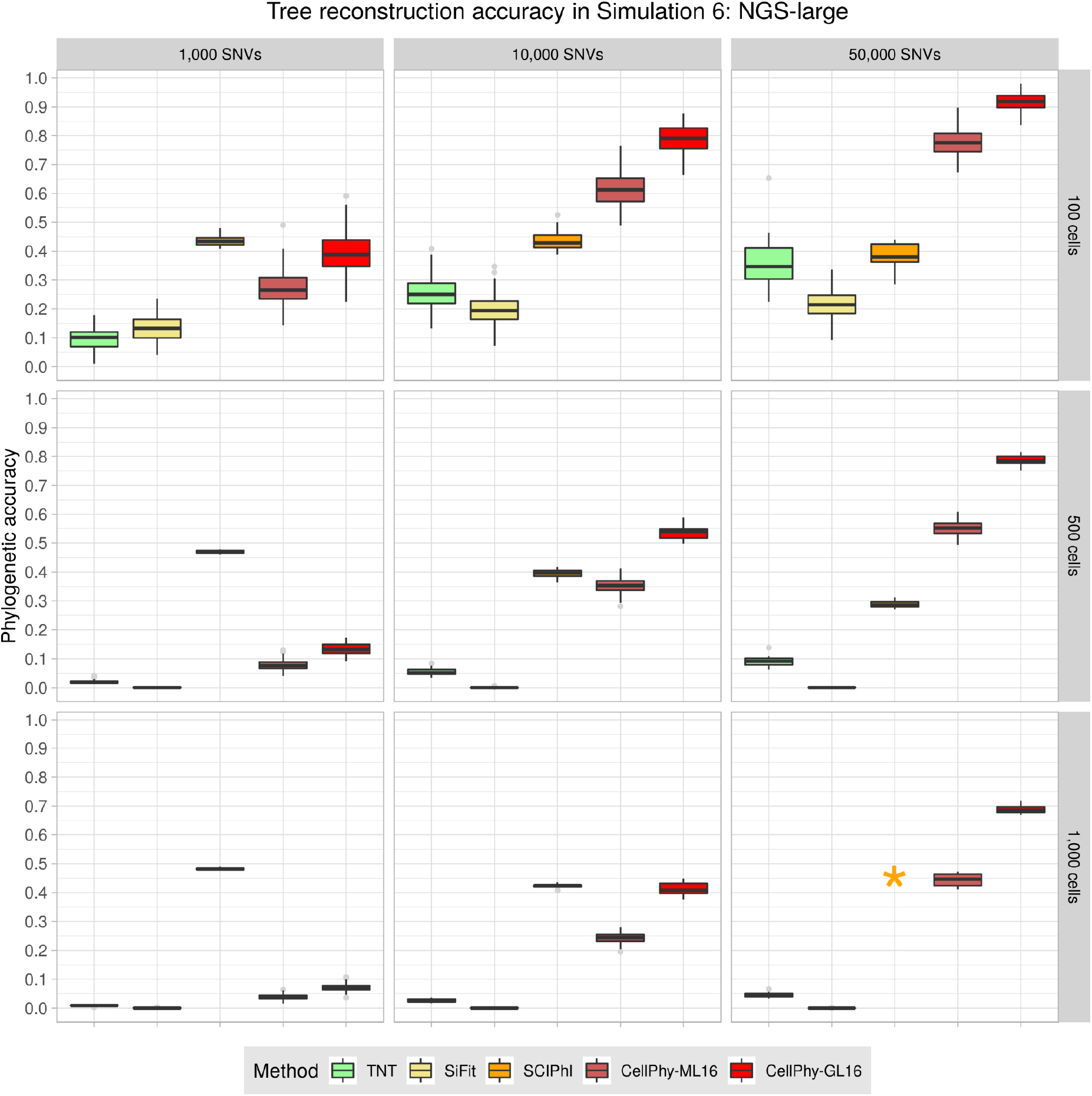
Phylogenetic reconstruction accuracy in Simulation 6 (“NGS-large”). Mutations were introduced according to signature S1, and the sequencing depth was 5x. Read counts were simulated with a 5% amplification error, 1% sequencing error, and 10% ADO. TNT, SiFiT, and CellPhy-ML16 use the inferred genotypes, CellPhy-GL16, uses the genotype likelihoods, and SCIPhI uses read counts. Phylogenetic accuracy is 1 – nRF (see Methods). For more than 1000 SNVs or more than 100 cells, only ten replicates were run for SiFit for reasonable running times. The orange star indicates that we could not obtain SciPhI results for the largest dataset (1,000 cells and 50,000 SNVs) after running the program for 100 hours. CellPhy was run using the ML genotypes error model (CellPhy-ML) and genotype likelihoods (CellPhy-GL). See Figure 1 for an explanation of the boxplots.

#### Estimation of genotype error and ADO rates

Besides inferring the ML tree, CellPhy can calculate ML estimates for the genotyping error and the ADO rate of scDNA-seq datasets. Across the different simulation scenarios described above, CellPhy estimated the genotyping error quite accurately (MSE: 0.00003 - 0.002), with a slight over/under estimation when its actual value was below/above 5%, respectively (Figure S8A). The ML estimates of ADO were more variable and tended to underestimate the true value (MSE = 0.002 - 0.02), but were still generally accurate, particularly at higher rates (Figure S8B). As expected, in both cases, improved estimates were obtained with larger datasets comprising more SNVs.

#### Computational speed

We compared the speed of the different methods by recording the running times for six simulated and two empirical data sets (Figure 5). TNT was the fastest method by at least two orders of magnitude, which is not surprising as parsimony scores are substantially cheaper to compute than likelihood scores. After TNT, CellPhy (under both ML and GL models) was the second-fastest method, being one to two orders of magnitude faster than SiFit, SCIPhI, infSCITE, or OncoNem. For some of the most extensive data sets, including simulated and empirical data, infSCITE and OncoNem did not finish after several days.

**Figure 5.**
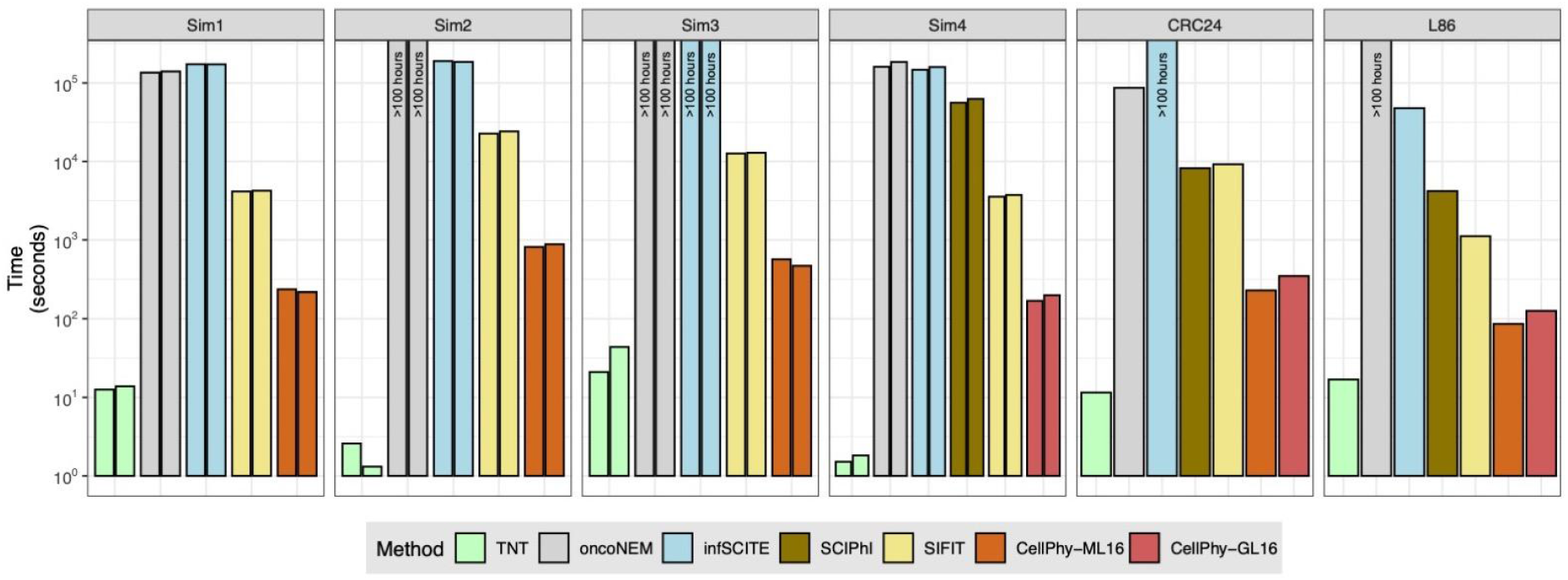
Speed comparisons for simulated and real datasets. “Sim1” corresponds to two simulated single-cell datasets with 40 cells and 4753 and 4761 SNVs. “Sim2” corresponds to two simulated datasets with 100 cells and 9935 and 9942 SNVs. “Sim3” corresponds to two simulated datasets with 60 cells and 9982 and 9985 SNVs. “Sim4” corresponds to two simulated datasets with 40 cells and 3986 and 4270 SNVs. CRC24 and L86 correspond to two empirical datasets (see Methods). Note the logarithmic time scale on the y-axis.

### Application to single-cell data

#### Phylogenetic reconstruction of a colorectal cancer

We analyzed a single-cell WGS dataset (CRC24) produced in our lab, consisting of 24 cells collected from two primary tumor biopsies of a patient with colorectal cancer (CRC). After filtering out germline polymorphisms, SNVs in non-diploid regions, and low-quality variants (see *Methods*), we identified a total of 18,099 SNVs. Some SNVs occurred in established CRC driver genes, such as *APC, BRAF, BRCA2, LRP1B*, and *MAP2K4*. In the ML tree estimated by CellPhy using the genotype likelihoods (“CellPhy-GL” model) (Figure 6A), cells tended to group according to their geographical location and phenotype, although not in a perfect fashion. Some of these relationships are well supported by the data, as reflected by several high bootstrap values, but others are not. This heterogeneous support illustrates one of the convenient features of CellPhy: it can provide phylogenetic confidence measurements for different parts of the tree. Interestingly, non-stem cells had, in general, longer branch lengths than stem cells, suggesting potential differences in the evolutionary rate between these two cell types. We mapped the non-synonymous mutations onto the internal branches of the tree using a custom script (see *Methods*). We found that all tumor cells share the vast majority of these mutations (i.e., they are clonal), including variants affecting genes previously associated with CRC progression (e.g., *INPP1, CDC5L, ROR2, EXOSC5*) (Li et al. 2000; Ma et al. 2016; Li et al. 2017; Pan et al. 2019). The tree topologies inferred by SiFit, SCIPhI, infSCITE, and TNT (Figure S9A-D) were distinct from the topology inferred by CellPhy (nRF=0.45, 0.63, 0.59, and 0.86, respectively), but with a similar, albeit not identical, overall pattern regarding geography and cell type. Unfortunately, for the SiFit, SCIPhI, and infSCITE trees, the absence of branch support measures makes it impossible to determine which parts of the estimated trees can be trusted.

**Figure 6.**
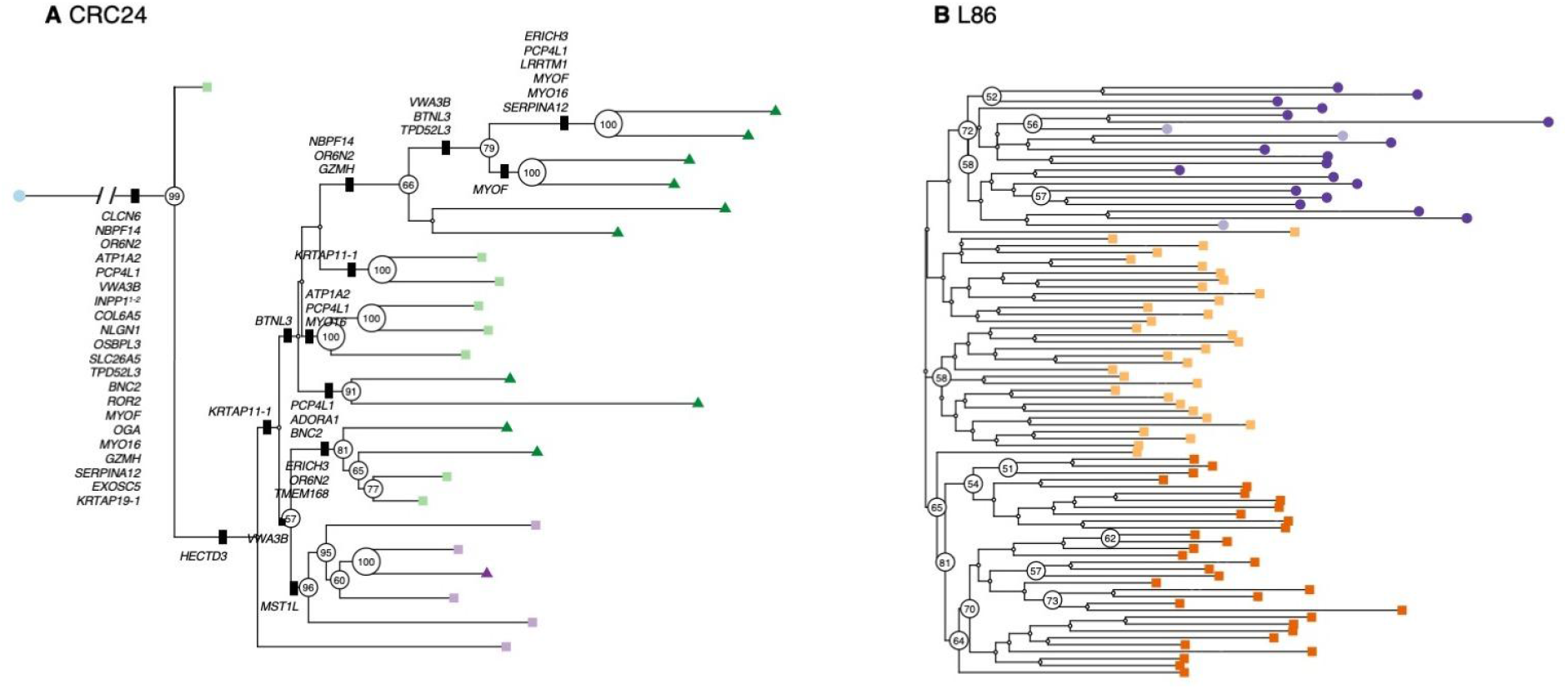
CellPhy tree for the CRC24 and L86 datasets. **(A)** “CellPhy-GL” CRC24 tree. Distinct shapes and colors represent cell type: healthy (blue circle); tumor-non-stem from TI region (dark green triangle), tumor-stem from TI region (light green square), tumor-non-stem from TM region (dark purple triangle), tumor-stem from TM region (light purple square). Only bootstrap support values above 50 are shown. Non-synonymous mutations are displayed on internal branches. **(B)** “CellPhy-GL” L86 tree. Distinct shapes and colors represent cell type: healthy diploid cells - from both primary and metastatic sites - (dark purple circle), healthy diploid cells missorted (light purple circle), primary tumor aneuploid cells (light orange square), metastatic aneuploid cells (dark orange square). Only bootstrap values above 50 are shown.

#### Revisiting the evolutionary history of a metastatic colorectal cancer

We also explored a published dataset from a metastatic colorectal cancer patient (Leung et al. 2017). In the original study, after performing custom targeted sequencing of 186 single-cells sampled from primary and metastatic lesions, the authors derived a cell tree using SCITE. They inferred a polyclonal seeding of liver metastases (i.e., distinct populations of tumor cells migrated from the primary tumor towards the liver). However, their findings have been recently re-evaluated in two different studies. Zafar et al. (2019) used the newly developed SiCloneFit, which relaxes the infinite-sites assumption, and proposed a polyclonal seeding of the metastases. More recently, Satas et al. (2020) performed a joint analysis of SNVs and copy-number variants (CNVs) with SCARLET and concluded instead that a single clone seeded the liver metastasis. As our analysis focuses on cancer history, to speed up computation, we removed the majority of the healthy cells in the original dataset, ending up with 86 cells (L86 data set). Using SC-Caller (Dong et al. 2017), we identified 597 SNVs distributed over the genome portion not affected by CNVs, including most variants detected in the original study (e.g., APC, *NRAS, MYH11, LINGO2, IL7R, F8, FUS*). In the CellPhy tree, all metastatic cells clustered together with high support, indicating a monoclonal origin of the liver metastasis (Figure 6B). Interestingly, while some primary (PA-54 and PA-27) and metastatic aneuploid (MA-25 and MA-26) cells appear intermixed with healthy diploid cells, these correspond to cells mislabelled during FACS sorting, as previously noted by the authors.

Furthermore, after mapping the non-synonymous mutations onto the internal branches of the CellPhy tree (Figure S10), we found that all cancer cells harbor somatic variants affecting genes that can contribute to human intestinal neoplasia (i.e., *MYH11* and *STAG1*) (Alhopuro et al. 2008; Romero-Pérez et al. 2019). Apart from CellPhy, only SCIPhI recovered a single metastatic clade (Figure S11). Although in the SiFit tree most metastatic cells cluster together, some of them appear intermixed with the primary tumor cells. In the infSCITE and TNT trees, the tumor cells did not form a clade, a result that does not appear to be very realistic.

#### Applicability of CellPhy to non-cancer data and bulk clonal sequences

Finally, we used CellPhy to analyze two non-cancer WGS single-cell datasets. The first of these datasets (E15) consists of 242 somatic SNVs from 15 single neurons from a healthy individual (Figure S12A) (Evrony et al. 2015). In the CellPhy tree built using genotype likelihoods (Figure S12A), different lineages seem to exhibit highly different evolutionary rates. However, most branches had low bootstrap support (<50%), suggesting that more SNVs are required before reliable interpretations can be made. All other methods recovered very distinct trees (Figure S12B-D), and in the case of TNT (SCIPhI, infSCITE, and SiFit do not assess branch support) also with shallow bootstrap values.

The second dataset (LS140) consists of 140 single cell-derived human hematopoietic stem and progenitor colonies from a healthy individual (Lee-Six et al. 2018). In this case, because there is no amplification involved, we expect a minimal genotype error and no ADO. Remarkably, CellPhy ML estimates for this data set were zero for both error and ADO parameters. The CellPhy tree shows very high bootstrap values (Figure S13), highlighting the quality of this dataset, which has a strong phylogenetic signal. We somehow expected this result given that the dataset has 127,884 SNVs and lacks single-cell biases. Moreover, our analysis confirms an early, well-supported divergence of the cell colonies into two distinct groups, together with a lack of geographical structure, reinforcing the idea of a continuous redistribution of stem cell pools at the whole-body level.

## Discussion

We have developed CellPhy, a phylogenetic tool for analyzing single-cell SNV data inspired by existing models and methods in statistical phylogenetics. Unlike some of its competitors (OncoNEM, infSCITE, and SCIPhI), CellPhy does not assume an infinite-site model of evolution, a widespread postulate in somatic evolution challenged by recent studies (Kuipers, Jahn, Raphael, et al. 2017; Demeulemeester et al. 2021). CellPhy’s evolutionary model explicitly considers all 16 possible phased DNA genotypes –but can work with both phased and unphased data– and can also account for their uncertainty by using genotype likelihoods as input. Furthermore, CellPhy can directly estimate single-cell error and ADO rates from the genotypes if the genotype likelihoods are unavailable. Finally, CellPhy can reconstruct ancestral states, predict mutations on tree branches, and provide a statistical measure of branch support. To benchmark CellPhy, we conducted computer simulations under different scenarios with varying degrees of complexity. Unlike in previous comparisons (Jahn et al. 2016; Ross and Markowetz 2016; Kuipers, Jahn, Raphael, et al. 2017; Zafar et al. 2017), we consider more realistic somatic genealogies by sampling sets of cells from a growing population. Demographic growth results in more difficult-to-reconstruct phylogenies, with shorter internal and longer terminal branches that require hundreds of SNVs to be accurately inferred. Overall, CellPhy was the most accurate method, both under infinite- and finite-site mutation models with different mutational patterns (e.g., using COSMIC trinucleotide mutational signatures), mainly when genotype errors and ADO were common, which is the case for scDNA-seq data. Our simulations suggest that accounting for SNV calling uncertainty is essential when sequencing depth is low to moderate, which is typically the case for single-cell WGS due to the sequencing costs. With a 5x sequencing depth, the ability to account for genotype uncertainty makes CellPhy substantially more accurate than its competitors. Notably, the accuracy of CellPhy does not come at the cost of speed. CellPhy is one to two orders of magnitude faster than SiFit, infSCITE, SCIPhI, or OncoNEM. Although, as expected, parsimony-based TNT was by far the fastest tool, this comes at the cost of considerably worse accuracy under most scenarios.

Our analysis of empirical scNDA-seq data suggests that CellPhy can provide biological insights where other methods fail to do so, with a seemingly more reliable grouping of cells according to phenotype or location. For example, in our re-analysis of the dataset from Leung *et al*. (2017), CellPhy was able to recover a monoclonal metastasis – in line with a recent analysis by Satas *et al. (2020)* that accounts for copy number variants–, while most competing methods implied a polyclonal seeding. Moreover, the CellPhy bootstrap analyses illustrate the importance of explicitly considering phylogenetic uncertainty. Without such a measure, one cannot assess if the data support the estimated phylogeny. Further, the analysis of cell colonies shows that CellPhy can also be used to estimate trees from clonal sequences that do not necessarily correspond to amplified single-cells.

Overall, our results suggest that CellPhy constitutes an efficient tool for estimating phylogenetic trees from scDNA-seq data. It is important to note that to achieve high phylogenetic accuracy under realistic conditions (i.e., growing populations with long terminal branches), CellPhy requires a reasonably high number of SNVs (hundreds to tens of thousands, depending on the number of cells). However, as can be seen in our benchmark, this limitation is common to all methods. It reflects the fundamental problem of poor signal-to-noise ratio: the faster the cell population grows, the more cells we have, and the higher the single-cell error and ADO rates are, the more SNVs are needed to recover the true cell relationships. We expect that further improvements in single-cell sequencing accuracy, variant calling sensitivity, and genotype phasing will yield datasets that are better suited for phylogenetic inference. Furthermore, some common simplifying assumptions of the classical phylogenetic models (reversibility, stationarity, context-independence) could be particularly problematic in the context of somatic evolution, which takes place at a much shorter temporal scale. Albeit methodologically and computationally challenging, eliminating those assumptions could improve accuracy on scDNA-seq datasets with a weak phylogenetic signal.

## Methods

### The CellPhy model

#### Model of nucleotide substitution for phased/unphased diploid genotypes

We developed a substitution model for diploid genotypes akin to those typically used for DNA sequences in organismal phylogenetics (see Felsenstein 2004). Specifically, we built a finite-site, continuous-time, Markov model of genotype evolution considering all possible 16 phased diploid states (A|A, A|C, …, T|T), in which SNVs are independent and identically distributed. This model, named GT16, is defined by a rate matrix Q that contains the instantaneous transition rates q*X*↔*Y* among genotypes *X* and *Y*. For computational convenience, we assume a time-reversible process, in which case the Q matrix is the product of a symmetric exchangeability matrix R (r*X*↔*Y* = r*Y*↔*X*) and a diagonal matrix of stationary genotype frequencies π_X_:

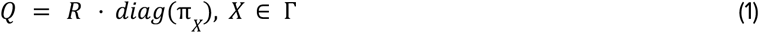

where Γ= {A|A, A|C, A|G, A|T, C|A, C|C, C|G, C|T, G|A, G|C, G|G, G|T, T|A, T|C, T|G, T|T}. We assume that only one of the two alleles in a genotype can change in an infinitesimal amount of time. Furthermore, because maternal and paternal chromosomes evolve independently in somatic cells, we also assume that the instantaneous transition rate of a given allele *a* in genotype *X* to allele *b* in genotype *Y* does not depend on the identity of the homologous allele *n* within the respective genotypes. In other words, we assume r(*na*↔*nb*) = r(*b↔a*). Conveniently, these assumptions significantly reduce the number of free parameters in the Q matrix, as we only need to consider five nucleotide exchangeabilities (*α* = r(*A↔C*), *β* = r(*A↔G*), *γ*= r(*A↔T*), *κ* = r(*C↔G*), λ = r(*C↔T*); let *μ* = r(*G↔T*) = 1) and 15 stationary phased genotype frequencies (π_A|A_, π_A|C_, π_A|G_, π_A|T_, π_C|A_, π_C|C_, π_C|G_, π_C|T_, π_G|A_, π_G|C_, π_G|G_, π_G|T_, π_T|A_, π_T|C_, π_T|G_; π_T|T_= 1 - ∑π_X_):

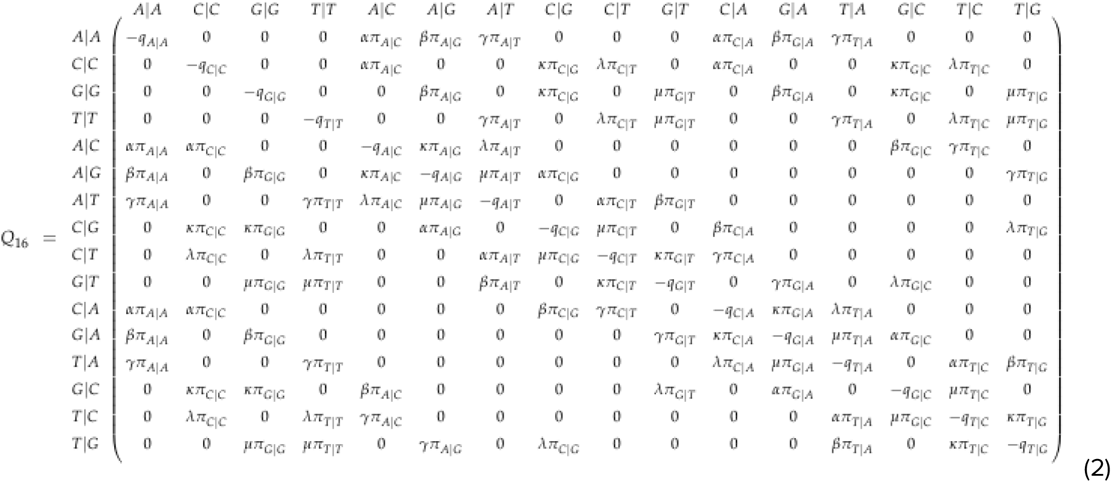

The probabilities of changing from a given genotype to another along a branch of length *t* are then given by:

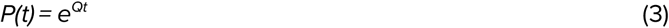

Notably, the GT16 model can also work with unphased genotype data simply by assigning the same relative likelihood at the tips of the tree to the two possible resolutions of the phase for the observed genotype (see Table S2). This flexibility is relevant because current scDNA-seq techniques do not reveal the phase of the genotypes (i.e., we do not know which allele is located in the maternal or paternal chromosome). Therefore, in our simulations (see below), the simulated genotypes will always be unphased.

In addition to the GT16 model, we also implemented a model for unphased genotypes with only ten states, called GT10 (see Supplementary Note 2 for details). The GT10 model is computationally less expensive, approximately twice as fast as GT16 (see Figure 5). To maintain the reversibility assumption -otherwise, the calculations are much more complex– the GT10 model assumes that the probability of change between homozygous and heterozygous genotypes is equivalent. However, this is not correct, as the change from homozygote to heterozygote is twice as likely as the change from heterozygote to homozygote. Despite this theoretical flaw, GT10 and GT16 showed very similar tree inference accuracy in our experiments.

#### Single-cell genotype error model

We assume that the true genotypes are always diploid and biallelic. To incorporate errors in the observed genotypes arising during single-cell whole-genome amplification (scWGA) and sequencing, we consider two free parameters: the allelic dropout (ADO) rate (*δ*) and the amplification/sequencing error (ERR) rate (*ε*).

Allele dropout occurs during scWGA when one of the two alleles is not amplified and is absent from the sequencing library. ADO always implies a single allele in our model, as if both alleles drop, no reads would be available, and no genotype could be observed (i.e., it would be “missing”). Therefore, *δ* is the probability that the amplification of one or the other allele has failed, and thus that we observe the homozygous genotype defined by the amplified allele. Given the phased genotype *a*|*b*, and with “_ “ indicating the dropped allele, we define the ADO rate as:

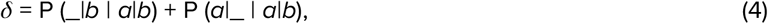

where P (*Y* | *X*) is the probability of observing genotype *Y* after sequencing, given the true genotype *X*.

Genotype errors other than ADO can result from polymerase errors during scWGA, in the course of sequencing, variant calling, or generally represent an incorrect allele. Importantly, we assume a maximum of one ERR per genotype. Given that ε tends to be small, in the order of 10^−3^ to 10^−5^ (Huang et al. 2015), the probability of two ERR in one genotype, ε^2^, is negligible. Specifically, *ε* is the probability that allele *a* will be observed as another allele *b* ≠ *a*:

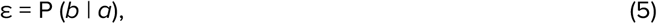

where *b* != *a*. Note that we allow for the presence of both ADO and ERR in the same observed genotype. Under these assumptions, if the true genotype is homozygous *a*|*a*, for phased genotypes P (*Y* | *X*) becomes:

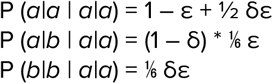

Likewise, if the true genotype is heterozygous *a|b*, P (*Y* | *X*) can be shown to be:

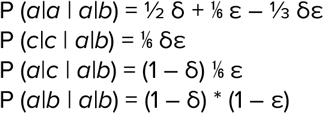

For the remaining scenarios, given the assumptions of the error model, P (*Y* | *X*) is zero. See Supplementary Material Note 1 for a detailed explanation.

Thus, the likelihood of the true genotype *X* for the SNV *i* and cell *j*, given the observed genotype *Y*, becomes:

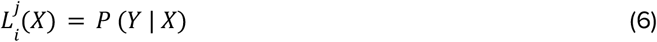

For missing data, we consider all possible genotypes to be equally likely, which is a standard assumption in likelihood-based phylogenetic inference. In this case:

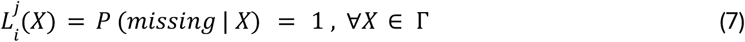

#### Phylogenetic likelihood

We consider the evolutionary history of cells as an unrooted binary tree, where *τ* is a tree topology and *t* is a vector of branch lengths. A branch length represents the mean number of expected mutations per SNV. We define the phylogenetic likelihood of the cell tree *T* as the conditional probability of the observed SNV matrix S given the substitution model *M* with parameters *θ* and *T*:

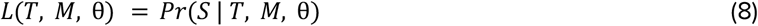

We assume that genomic sites evolve independently under model *M*. Therefore, the probability of observing the SNV matrix *S* is the product over the independent probabilities for all individual SNVs, *S_i_*:

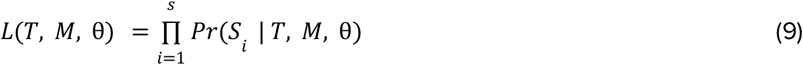

where *s* is the total number of SNVs. For numerical reasons, that is, to avoid floating-point underflow, in practice, we calculate the log-likelihood score:

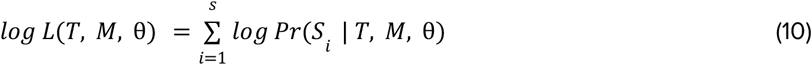

Let us place an additional imaginary node, called *virtual root*, into an arbitrary branch of the unrooted tree *T* at a random position along that branch. Without loss of generality, if we assume that this virtual root node has an index of 0, we can index the tip nodes from 1 to *c* by their respective cell numbers and index the internal nodes from *c*+1 to 2*c*–2 (note that an unrooted binary tree with *c* leaves has *c*–3 inner nodes, i.e., (2*c*–2) – (*c*+1) = *c*–3). Then, under the GT16 model, per-SNV probabilities can be computed as follows:

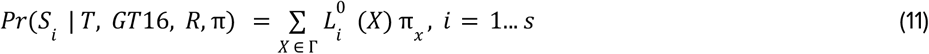

where *X* is a genotype, π_*x*_ is the stationary frequency of the genotype *X*, and 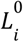 is the vector of genotype likelihoods at the virtual root, which we compute as follows:

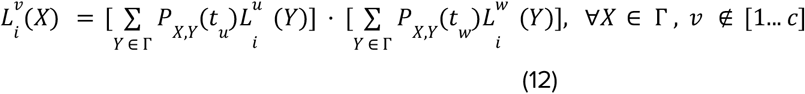

where *Y* is another genotype, *u* and *w* are both children nodes of *v* in the direction from the virtual root (Figure S14) for the respective branch lengths.

We initialize the genotype likelihood vectors at the tip nodes 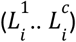 depending on the input type (see Section “Input data” below) and error model. If the input is the genotype matrix *S*, in the absence of observational error, these tip likelihood vectors are:

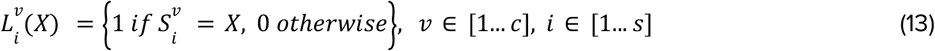

While if we include the error model, the tip likelihoods are computed according to the equations given in the “Single-cell genotype error model” section.

Otherwise, if the input consists of genotype likelihoods *G*, the tip likelihoods are:

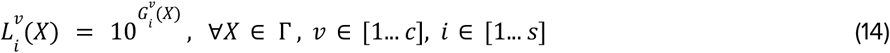

Note that the GT16 model will consider both phasing options equally likely when the input genotype likelihoods are unphased.

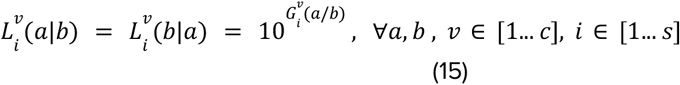

where *a* and *b* represent any two alleles.

Finally, if Phred-scaled likelihoods for REF/REF, REF/ALT, and ALT/ALT genotypes are provided (PL field in a VCF file), we calculate tip likelihoods as follows:

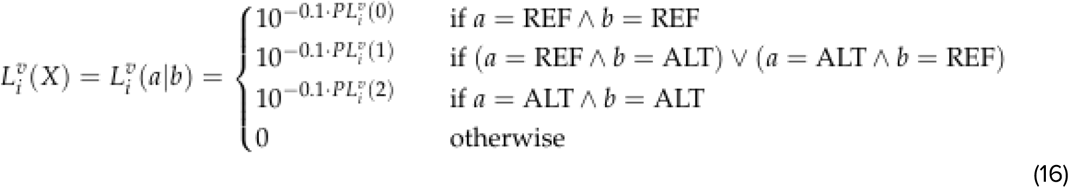

The final tree can be rooted using an outgroup (see Felsenstein 2004). For the sake of completeness, we included a simple explanation of standard phylogenetic likelihood calculations on DNA sequence alignments in the Supplementary Material (Figures S14-S16, Supplementary Note 3).

### Implementation

#### Overview

We implemented CellPhy as a pipeline based on a modified version of RAxML-NG (Kozlov et al. 2019). In addition to the core tree search functionality of RAxML-NG, the CellPhy pipeline offers features such as VCF conversion, mutation mapping, and tree visualization using the *ape* (Paradis and Schliep 2019) and the *ggtree* package (Yu et al. 2017). Furthermore, CellPhy provides reasonable defaults for most parameters, which allows the user to run a “standard” CellPhy analysis by specifying just the input VCF file (or the genotype matrix). Alternatively, expert users can customize every aspect of the CellPhy analysis to fit their needs, as we show in the tutorial (https://github.com/amkozlov/cellphy/blob/master/doc/CellPhy-Tutorial.pdf). In the remainder of this section, we will also provide implementation details on each step of the CellPhy pipeline. CellPhy code and documentation are freely available at https://github.com/amkozlov/cellphy.

#### Input data

CellPhy accepts two types of input, a matrix of genotypes in FASTA or PHYLIP format or a standard VCF file (https://samtools.github.io/hts-specs/VCFv4.3.pdf). When the input is a genotype matrix, genotypes are encoded as shown in Table S2. When the input is a VCF, CellPhy can run in two distinct modes. The first mode (“CellPhy-ML”) requires a VCF with at least the GT field (that stores the genotype calls), in which case CellPhy simply extracts the genotype matrix. The second mode (“CellPhy-GL”) requires a VCF with the PL field (which stores the Phred-scaled genotype likelihoods) and uses the likelihood of each genotype instead. While commonly used variant callers for single-cell data (e.g., Monovar (Zafar et al. 2016)) generate VCF files with a standard PL field, users should be aware that the PL definition may differ from its standard meaning in different callers. Indeed, SC-Caller (Dong et al. 2017), for instance, uses the PL field not only to store the likelihood of heterozygous and alternative homozygous genotypes but also the likelihood of sequencing noise and amplification artifacts. On this basis, the PL field in VCF files stemming from SC-Caller needs to be converted to the standard PL format before CellPhy can be used (see Table S3 outlining CellPhy’s compatibility with popular variant calling algorithms).

#### Phylogenetic tree search

CellPhy uses the broadly used and well-tested heuristic tree search strategy of RAxML (Stamatakis 2014) and RAxML-NG (Kozlov et al. 2019). CellPhy’s search algorithm starts by default with 20 different trees, ten obtained using a maximum parsimony-based randomized stepwise addition order routine and 10 with a completely random topology. The ML tree search itself alternates between model parameter optimization and tree topology optimization phases. The key mechanism for searching tree topologies are so-called subtree pruning and regrafting (SPR) (Robinson 1971) moves that attempt to remove subtrees from the tree and subsequently place them into different branches to assess if the likelihood improves. Those SPR moves are applied iteratively until no SPR move further increases the tree’s likelihood. In this case, CellPhy terminates and returns the best-found ML tree. For further details, please see (Kozlov 2018: 4.2.3) and references therein.

#### Model parameter optimization

CellPhy uses the L-BFGS-B method (Fletcher 2000) to optimize genotype substitution rates and equilibrium base frequencies. ADO rate and amplification/sequencing error rate are optimized independently using Brent’s method (Brent 1971). After each iteration of Brent’s algorithm, CellPhy re-computes all per-genotype likelihoods according to Eq. 7 using the new values of the ADO rate *δ*’ and the amplification/sequencing error ε’.

#### Branch support

CellPhy can compute confidence values for individual inner branches of the ML tree using two bootstrap (BS) techniques, the standard BS (Felsenstein 1985) and the transfer BS (Lemoine et al. 2018). In the standard BS, the first step consists of generating many BS replicates, typically 100 to 1,000, from the original dataset by randomly sampling SNV sites with replacement. Then, an ML tree is estimated for each replicate. Finally, the support for each inner branch in the ML tree is computed as the percentage of BS trees that contain that branch, as outlined in the example provided in Figure S16. The standard BS only considers exact matches (i.e., the branch in the ML tree and the BS trees must match exactly to be counted). In contrast, the transfer BS also considers inexact matches to account for the tree search uncertainty and vastness of the tree search space in phylogenetic analyses with many cells.

#### Mapping mutations onto the tree

CellPhy can show predicted mutations on the branches of the inferred cell tree. To this end, it performs marginal ancestral state reconstruction (Yang et al. 1995) to obtain the ML genotype for every SNV at every inner node of the tree. At the tips of the tree, occupied by the observed cell genotypes, depending on the input, CellPhy applies Eq. 4 to compute genotype likelihoods given the observed genotype *y* and estimated error rates (*δ*, *ε*) or directly uses the genotype likelihoods provided in the VCF file. Then, it compares ML genotypes between two nodes connected by a branch, and if they differ, a mutation is predicted on the corresponding branch. The mutation mapping output consists of two files, a branch-labeled tree in the Newick format and a text file with a list of predicted mutations (SNV names or positions) at each branch. We also provide a script (https://github.com/amkozlov/cellphy/blob/master/script/mutation-map.R) that automatically generates a plot with the mutations mapped onto the resulting phylogenetic tree, together with a tutorial that explains its use.

#### Computational efficiency

RAxML-NG was developed with a particular focus on high performance and scalability to large datasets. Hence, CellPhy capitalizes on numerous computational optimizations implemented therein, including highly efficient and vectorized likelihood calculation code, coarse- and fine-grained parallelization with multi-threading, checkpointing, and fast transfer bootstrap computation (Lutteropp et al. 2020).

### Benchmarking

We used computer simulations to benchmark the accuracy of CellPhy under different scenarios (Table S1), relative to the state-of-the-art methods for single-cell phylogenies OncoNem (Ross and Markowetz 2016), infSCITE (Jahn et al. 2016; Kuipers, Jahn, Raphael, et al. 2017), SiFit (Zafar et al. 2017), and SCIPhI (Singer et al. 2018). For the first three methods, the input is the matrix of observed reference/non-reference homozygous/heterozygous genotypes. For SCIPhI, the input is a mpileup file with site read counts. Also, we included in the comparison the standard phylogenetic method TNT (Goloboff and Catalano 2016). TNT implements a maximum parsimony (MP) approach and therefore attempts to find the tree/s that require the least number of mutations to explain the data. TNT is very popular in MP organismal phylogenetics, heavily optimized for computational speed and efficiency. It is not designed for single-cell NGS data and therefore assumes that the observed genotypes are error-free.

#### Simulation of genealogies and genotypes

For the simulation of single-cell diploid genotypes, we used CellCoal (Posada 2020). This program can simulate the evolution of a set of cells sampled from a growing population, introducing single nucleotide variants on the coalescent genealogy under different models of DNA mutation. Furthermore, it can also introduce the typical errors of single-cell sequencing, specifically ADO, amplification, and sequencing errors, and doublets, either to the observed genotypes or directly into the read counts (Fig. S1).

We designed six distinct simulation scenarios (Simulation 1 - 6) representing different types of scDNA-seq datasets (Table S1), including variable numbers of cells (40-1000) and sites (1000-50000). We simulated unphased genotype data in all cases, as current scDNA-seq techniques do not reveal the genotype phase. We chose a set of scenarios and parameter values that, in our opinion, are representative of different situations that researchers are likely to encounter. The cell samples were assumed to come from an exponentially growing population (growth rate equal to 1×10^-4^) with a present-day effective population size of 10000. Across scenarios, we set a constant value of 0.1 for the root branch. Note that the mutations in this branch are shared by all cells (Fig. S1). We also defined an outgroup branch length (Fig. S1) of zero in all cases, so the healthy cell and the most recent common ancestor (MRCA) of the sample (a single healthy cell plus several tumor cells) have identical genotypes. The standard coalescent process results in an ultrametric tree, where all tips have the same distance from the MRCA of the sample. However, we introduced rate variation across cell lineages by multiplying the branch lengths of the resulting coalescent genealogy with scaling factors sampled from a Gamma distribution with a mean of 1.0 (see Yang 1996).

Only in the first simulation scenario, we considered a fixed number of SNVs. In the remaining scenarios, the number of observed mutations resulted from applying a mutation rate of 1×10^−6^ (Martincorena and Campbell 2015), plus the different scDNA-seq errors. We explored different infinite- and finite-sites mutation models at the single nucleotide or trinucleotide level. Except for the ISM scenarios (Simulations 1 and 3), we introduced a variation of the mutation rate across sites using a Gamma distribution with a mean of 1.0 (as in Yang 1996).

We simulated unphased genotypes, as current scDNA-seq techniques do not reveal the phase. We generated the observed genotype matrices in two distinct ways. In the first three scenarios (Simulations 1-3), we obtained the observed genotypes by directly adding sequencing/amplification errors (i.e., changing one or both alleles) and ADO to the simulated genotype matrices. In the other three (Simulations 4-6), the generation of the observed genotypes was more complex. In this case, we first simulated read counts for each cell based on the true genotypes, considering different overdispersed sequencing depths, as well as amplification and sequencing errors. For simplicity, we assumed that maternal and paternal chromosomes are amplified with the same probability. Also, we consider that the number of reads is half for those genomic positions in which only one allele was amplified. In Simulation 5, we introduced so-called doublets, that is, two cells that are erroneously sequenced together and that thus appear as a single cell in the sequencing data. For every combination of parameters, we generated 100 replicates. In total, we generated 19,400 cell samples.

##### Simulation 1: infinite-site model, low number of SNVs (“target-ISM”)

We started with a simple scenario with 40 sampled cells and 250, 500, or 1000 SNVs, assuming a diploid ISM model (Kimura 1969). Under this model, a given site can only accumulate a single mutation along with the genealogy, either in the maternal or paternal chromosome. We introduced genotype errors and ADO at different rates (Table 1).

##### Simulation 2: finite-site model, large number of SNVs (“WGS-FSM”)

In this case, we simulated a larger number of SNVs, more typical of whole-genome sequencing (WGS) experiments. The number of sampled cells was 100. The mutation model, in this case, was a non-reversible version of the finite-site General Time Reversible Markov model (Tavaré 1986), that we called GTnR, assuming a set of single-nucleotide instantaneous rates extrapolated (essentially, we pooled the same mutation in the same rate independently of the 5’ and 3’ context) from the trinucleotide mutational signature 1 at COSMIC (https://cancer.sanger.ac.uk/cosmic/signatures):

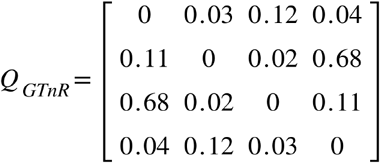

The overall mutation rate was set to 1×10^−6^, which resulted in about 2000 *true* SNVs (see Table 1). However, since ADO events and genotype errors can introduce false negatives and false positives, the number of *observed* SNVs ranged between 1147 and 10000. The mutation rates varied across sites according to a Gamma distribution (+G) with shape parameter and mean equal to 1.0 (i.e., moderate among-site rate heterogeneity).

##### Simulation 3: mutational signatures, large number of SNVs (“WGS-sig”)

This scenario is similar to the previous one, with 60 cells and assuming a trinucleotide ISM model, with COSMIC signatures 1 and 5. The former is a ubiquitous signature in human cells with a predominance of C>T transitions in the NCG trinucleotide context and related to the spontaneous deamination of 5-methylcytosine (Alexandrov et al. 2013). The latter is also a typical age-related signature with a predominance of T>C substitutions in the ATN trinucleotide context, related to transcriptional strand bias (Alexandrov et al. 2015).

##### Simulation 4: genotype likelihoods from NGS read counts (“NGS-like”)

In this scenario, and the next two, we simulated NGS read counts from the simulated genotypes. The number of sampled cells was 40, with 10000 sites and the same mutation model (GTnR+G) and mutation rates as in Simulation 3. We explored three sequencing depths (5x, 30x and 100x), three ADO rates (0, 0.05, 0.10), three amplification error rates (0, 0.05, 0.10), and three sequencing error rates (0, 0.01, 0.05). We assumed that amplification and sequencing errors among the four nucleotides were equally likely. From the read counts, CellCoal can also simulate the likelihood for all ten possible unphased genotypes at each SNV site, in this case under a 4-template amplification model (Posada 2020). The input for CellPhy was either the ten genotype likelihoods at each SNV site (CellPhy-GL mode) or the unphased genotype with the maximum likelihood (ML) value (CellPhy-ML mode). The input for SCIPhi was the read counts, while for the rest of the programs, it was the ML unphased genotype. In the rare case of tied ML genotypes, we chose one of them at random.

##### Simulation 5: NGS doublets (“NGS-doublet”)

In this case, we intended to explore the effects of doublets in the data. Settings were very similar to those for Simulation 4 but, for simplicity, we fixed the sequencing depth to 5x and explored two amplification error rates (0, 0.05), two sequencing error rates (0, 0.01), and four doublet rates (0, 0.05, 0.10, 0.20).

##### Simulation 6: NGS for large numbers of cells and SNVs (“NGS-large”)

Finally, to assess the scalability of the tools, we simulated scenarios with 100, 500, or 1000 cells and with 1000, 10000, or 50000 sites. Given the mutation rate, a large number of cells, and, most importantly, the amplification and sequencing error rates, almost all sites were observed as SNVs. Settings were very similar to those specified for Simulation 5 but, for simplicity, we fixed the sequencing depth to 5x and explored only one amplification (0.05) and one sequencing (0.01) error value. We only analyzed the first 20 replicates in this scenario due to the very high computational cost and prohibitive running times for several competing tools.

#### Settings for the phylogenetic analyses

##### Coding DNA into ternary genotypes

Our simulations produce unphased DNA genotypes with ten possible states. However, except for CellPhy, existing tools work with an alphabet composed of 0 (homozygous for the reference allele), 1 (heterozygous), 2 (homozygous for the alternative allele), and 3 (missing genotype). Therefore, we had to encode the simulated DNA genotypes into ternary genotypes (0-3). For this, we used the true reference allele, considering that, in real life, we usually know which allele is the reference. For the sake of simplicity, we did not introduce germline mutations. Importantly, our simulations do not necessarily produce bi-allelic SNVs, as in the finite-site model multiple mutations can coincide, and amplification and sequencing errors can also result in new alleles called. CellPhy does not have any limitation concerning the number of alleles at an SNV site, but competing tools handle multi-allelic sites differently. We explored three ways of coding sites with more than two alleles into ternary genotypes:

- Option *keep*: transform all heterozygotes to “1” and all homozygotes for the alternative allele/s to “2”. In this case, all simulated sites are held, regardless of the number of observed alleles.
- Option *remove*: eliminate sites from the data with more than two alleles. The final genotype matrix includes only bi-allelic sites.
- Option *missing*: keep only those genotypes that contain the reference allele and/or the major (most common) alternative allele. All other genotypes (containing minor alternative alleles) are considered as missing data (“3”). Therefore, the final ternary genotype matrix includes the same number of sites as the original DNA genotype matrix.

We considered all three encoding options only in Simulations 1 and 2. In the remaining simulations, we used only the ‘missing’ option, as it maximized accuracy in most cases.

##### TNT settings

We performed the TNT analyses using a binary data matrix in TNT format for all simulated and empirical datasets. We allowed 1000 trees to be retained for each run and performed tree searches by setting *mult = replic 100*. We stored all equally parsimonious trees and used additional ttags to store branch lengths and bootstrap support values.

##### OncoNEM settings

OncoNEM analyses were performed following the recommended settings in the OncoNEM vignette, but only for Simulation 1 due to its heavy computational requirements. We set the false positive rate as the actual genotype error for each scenario and performed a tree search for 200 iterations.

##### infSCITE settings

For Simulations 1-3, we ran infSCITE using a ternary data matrix composed of 0, 1, 2, and 3, as described above, and set the false positive rate (*-fd*) to be the actual genotype error for each simulated scenario. We set the false-positive rate for Simulations 4-5 as the sum of the simulated sequencing and amplification error rates. For the empirical analyses, we set the false positive rate to *1e-05*. We set the remaining parameters to the default values in all runs and obtained results after running an MCMC chain with 5 million steps, a fair trade-off between runtime and apparent MCMC convergence (the best tree score barely changed after 1M iterations).

##### SiFit settings

For all simulated and empirical datasets, we ran SiFit for 200000 iterations. The command line was: *java -jar /SiFit.jar -m <CELLS> -n <SNVS> -r 1 -iter 200000 -df 1 -ipMat snv_hap.XXX.sf -cellNames snv_hap.XXXX.names*

##### SCIPhI settings

Given that SCIPhI requires read counts to perform variant calling and phylogenetic reconstruction jointly, we ran it only for Simulations 4–6. We set the mean error rate (-wildMean) for each scenario as the sum of the true sequencing and amplification error rates. The command line was: *sciphi -o XXX.Result --in sampleNames -u 1 --ncf 0 --md 1 --mmw 4 --mnp 1 --ms 1 --mc 1 --unc true -l 200000 --seed $RANDOM XXX.mpileup --ese 0 --wildMean <ERROR RATE>*.

For the empirical datasets, to make the results comparable, we used samtools to generate mpileups for the positions previously identified by SC-Caller. These were in turn used as input for SCIPhI: *sciphi -o <SET>.SCIPhI --in <SET>-SampleNames.txt -u 0 --ncf 0 --md 0 --mmw 4 --mnp 1 --unc true -l 200000 --seed 421 <SET>.SCCaller-Positions.mpileup*

##### CellPhy settings

For Simulations 1-6, we performed a heuristic tree search starting from a single parsimony-based tree under the GT16 model. The command line for runs with ML genotypes as input was *cellphy.sh RAXML --search --msa snv_hap.XXXX.phy --model GT16+FO+E --tree pars{1}*. The command line for runs where the input were genotype likelihoods (VCF) was *cellphy.sh RAXML --search --msa vcf.XXXX --model GT16+FO --tree pars{1}*. For all empirical datasets except LS140, to take advantage of the genotype likelihood model, we used an *in-house* bash script (sc-caller-convert.sh, distributed together with CellPhy) to convert the PL field from our SC-Caller VCFs. In short, following SC-Caller authors’ suggestion, we used the highest likelihood score of the first two values in the PL field (i.e., sequencing noise, amplification artifact) as the Phred-scaled genotype likelihood of the reference homozygous (0/0) genotype, and the remaining values as the likelihood for heterozygous (0/1) and alternative homozygous (1/1) genotypes, respectively. Afterward, we ran CellPhy using the following command line *cellphy.sh RAXML --all --msa XXXX.vcf --model GT16+FO --bs-metric fbp,tbe --bs-trees 100* to perform an all-in-one analysis (ML tree inference and bootstrapping based on 100 bootstrap trees). For LS140, since we only had the genotype matrix available and given that these data were generated without whole-genome amplification, we ran CellPhy without the single-cell error model using the following command line *cellphy.sh RAXML --all --msa XXXX.vcf --model GT16+FO --prob-msa off --bs-metric fbp,tbe --bs-trees 100*.

#### Evaluation of phylogenetic accuracy

We defined *phylogenetic accuracy* as one minus the normalized Robinson-Foulds (nRF) distance (Robinson and Foulds 1981) between the inferred tree and the (true) simulated tree. This normalization consists of dividing the (absolute) RF distance by the total number of (internal) branches in *both* trees. Hence, the nRF distance is a convenient metric from zero to one that reflects the proportion of branches (bipartitions of the data) correctly inferred.

#### Running time comparisons

We characterized the computational efficiency of CellPhy by comparing running times for all methods on six datasets from Simulations 1-4 (sim1-ADO:0.50, ERR:0.10, sim2-ADO:0.10, ERR:0.05, sim3-ADO:0.15, ERR:0.10, Signature1, and sim4-Number of cells:100, ADO:0.25, Amp error:0.10, Seq error: 0.05) and two empirical datasets (CRC24 and L86) described below. We defined central processing unit (CPU) running time as the real-time returned by the Linux/Unix ‘time’ command. We ran all analyses on a single core from an Intel Xeon E5-2680 v3 Haswell Processor 2.5GHz with 128 Gb of RAM.

#### Analysis of empirical data

##### In-house single-cell WGS data from a colorectal cancer patient (CRC24)

We obtained a fresh frozen primary tumor and normal tissues from a colorectal cancer patient from the Biobank of I.D.I.S.-C.H.U.S. (PT13/0010/0068), part of the Spanish National Biobank Network. We processed the sample following the Ethical and Scientific Committee’s approval (CAEI Galicia 2014/015). We isolated EpCAM+ cells with a BD FACSAria III cytometer and amplified the genomes of 24 cells with Ampli1 (Silicon Biosystems). Moreover, we classified the isolated cells as stem or non-stem according to stemness markers at the cell surface (Stem: EpCAM+/Lgr5+/CD44-/CD166-; Non-stem: EpCAM+/Lgr5-/CD44-/CD166-). For each cell, we built whole-genome sequencing libraries using the KAPA (Kapa Biosystems) library kit. Each library was then sequenced at ^~^6x on an Illumina Novaseq 6000 at the National Center of Genomic Analysis (CNAG-CR; https://www.cnag.crg.eu/).

##### Retrieval of publicly available datasets (L86, E15, and LS140)

We also analyzed three public data sets with 86, 15, and 140 cells (referred to as L86, E15, and LS140, respectively). The L86 dataset consists of targeted sequencing data from 86 cells from a metastatic colorectal cancer patient (Leung et al. 2017) that we downloaded from the Sequence Read Archive (SRA) in FASTQ format, together with paired healthy-tumor bulk cell population samples (accession number: SRP074289). The E15 dataset consists of WGS data from 15 neurons (Evrony et al. 2015) from a healthy donor, downloaded from the SRA in FASTQ format, together with a bulk cell population from heart tissue (accession number: SRP041470). The LS140 dataset consists of 140 single cell-derived human hematopoietic stem and progenitor colonies from a healthy individual (Lee-Six et al. 2018). For this dataset, we directly downloaded the substitution calls from Mendeley data archive (https://data.mendeley.com/datasets/yzjw2stk7f/1).

##### NGS data processing and variant calling

We aligned single-cell and bulk reads to the human reference GRCh37 using the MEM algorithm in the BWA software (Li 2013). The mapped reads were then independently processed for all datasets by filtering reads displaying low mapping quality, performing local realignment around indels, and removing PCR duplicates. For the tumor bulk samples (i.e., CRC24 & L86), we obtained SNV calls using the paired-sample variant-calling approach implemented in the MuTect2 software (Cibulskis et al. 2013). For the E15 dataset, we ran HaplotypeCaller from the Genome Analysis Toolkit (GATK) (Poplin et al. 2018) software on the bulk sample from the heart tissue to identify and remove all germline variants.

In parallel, we used the single-cell SC-caller software (Dong et al. 2017) to retrieve single-cell SNV calls. In short, for each single-cell BAM we ran SC-Caller together with the corresponding healthy bulk DNA as input under default settings. Since different amplification methods were used to generate each dataset, we defined the bias estimation interval (--*lamb*) as the average amplicon size of each amplification method - 10,000 for MDA-based protocols (L86, E15) and 2,000 for Ampli1 (CRC24). Besides, since the actual genomic targets of the L86 dataset were not available, we ran SC-caller on the entire exome. We applied a series of heavy filters (see below) to remove potential off-target calls. We additionally estimated copy-number variants (CNVs) for each single-cell dataset. For the sc-WGS datasets (CRC24 and E15), we obtained CNV calls with the Ginkgo software (Garvin et al. 2015) using variable-length bins of around 500 kb. As for the L86 dataset, we determined CNVs using CNVPanelizer, an algorithm specifically designed to infer copy number states from targeted sequencing data.

We filtered our raw single-cell VCFs by excluding short indels, SNVs with a flag other than “True” in the SO format field (i.e., showing weak evidence of being a true somatic mutation), and variable sites with an alternative read count < 3. We also excluded variable sites in which the ML genotype estimate was above 120 (Phred-scaled). Such uncertainty in the genotype call was usually associated with sites experiencing an apparent disparity in the proportion of both alleles (i.e., allelic bias). Moreover, as we are primarily interested in analyzing diploid genomic regions, we removed those SNVs located within CN variable regions.

For each dataset, we then merged single-cell VCFs using the bcftools software (Li 2011) and applied a “consensus” filter to only retain sites present in at least one cell and the bulk tumor sample, or in two cells. For the E15 dataset, we limited this “consensus” filter to somatic sites observed in at least two cells, as we classified as germline all variants observed in the bulk sample. Finally, we removed positions missing (i.e., not covered by any read) in more than 50% of the cells and SNVs comprising more than one alternative allele. For the L86 dataset, we filtered out off-target SNVs located outside exonic regions. For the LS140 dataset, we converted the binary genotype matrix into a VCF by transforming 0, 1, and NA values into 0/0, 0/1, and ./., respectively. Afterward, we removed all duplicated (non-biallelic) positions and indels.

## Data availability

Raw single-cell whole-genome sequencing data from CRC24 have been deposited in the Sequence Read Archive (SRA - https://www.ncbi.nlm.nih.gov/sra) database under the accession code XXXXX. We have additionally analyzed previously published single-cell data sets (Evrony et al. 2015; Leung et al. 2017). Raw sequencing data for these sets are available from the SRA database under accession numbers SRP074289 (L86) and SRP041470 (E15). Furthermore, we generated the genotype matrix for the LS140 dataset (Lee-Six et al. 2018) from the substitution calls available at the Mendeley data archive (https://data.mendeley.com/datasets/yzjw2stk7f/1).

## Acknowledgments

We want to thank Debora Chantada, Pilar Alvariño, and Sonia Prado for their help in obtaining the CRC24 dataset and Phylogenomics lab members for their comments.

## Author contributions

A.K. implemented the model within RAxML-NG and ran part of the simulations. J.M.A. ran part of the simulations and performed the empirical analyses. A.S. supervised the phylogenetic implementation. D.P. conceived the CellPhy model, developed the error model, and defined the experimental design. All authors contributed to manuscript writing.

## References

Alexandrov LB, Jones PH, Wedge DC, Sale JE, Campbell PJ, Nik-Zainal S, Stratton MR. 2015. Clock-like mutational processes in human somatic cells. Nat. Genet. 47:1402–1407.

Alexandrov LB, Nik-Zainal S, Wedge DC, Aparicio SAJR, Behjati S, Biankin AV, Bignell GR, Bolli N, Borg A, Børresen-Dale A-L, et al. 2013. Signatures of mutational processes in human cancer. Nature 500:415–421.

Alhopuro P, Phichith D, Tuupanen S, Sammalkorpi H, Nybondas M, Saharinen J, Robinson JP, Yang Z, Chen L-Q, Orntoft T, et al. 2008. Unregulated smooth-muscle myosin in human intestinal neoplasia. Proc. Natl. Acad. Sci.U. S. A. 105:5513–5518.

Baslan T, Hicks J. 2017. Unravelling biology and shifting paradigms in cancer with single-cell sequencing. Nat. Rev.Cancer 17:557–569.

Brent RP. 1971. An algorithm with guaranteed convergence for finding a zero of a function. The Computer Journal 14:422–425.

Cibulskis K, Lawrence MS, Carter SL, Sivachenko A, Jaffe D, Sougnez C, Gabriel S, Meyerson M, Lander ES, Getz G. 2013. Sensitive detection of somatic point mutations in impure and heterogeneous cancer samples. Nature Biotechnology 31:213–219.

Demeulemeester J, Dentro SC, Gerstung M, Van Loo P. 2021. Biallelic mutations in cancer genomes reveal local mutational determinants. bioRxiv [Internet]. Available from: https://www.biorxiv.org/content/10.1101/2021.03.29.437407v1.abstract

Dong X, Zhang L, Milholland B, Lee M, Maslov AY, Wang T, Vijg J. 2017. Accurate identification of single-nucleotide variants in whole-genome-amplified single cells. Nature Methods 14:491–493.

Evrony GD, Lee E, Mehta BK, Benjamini Y, Johnson RM, Cai X, Yang L, Haseley P, Lehmann HS, Park PJ, et al. 2015. Cell lineage analysis in human brain using endogenous retroelements. Neuron 85:49–59.

Felsenstein J. 1981. Evolutionary trees from DNA sequences: a maximum likelihood approach. J. Mol. Evol. 17:368–376.

Felsenstein J. 1985. Confidence limits on phylogenies: an approach using the bootstrap. Evolution 39:783–791.

Felsenstein J. 2004. Inferring phylogenies. Sinauer associates Sunderland, MA

Fletcher R. 2000. Practical Methods of Optimization. Available from: http://dx.doi.org/10.1002/9781118723203

Garvin T, Aboukhalil R, Kendall J, Baslan T, Atwal GS, Hicks J, Wigler M, Schatz MC. 2015. Interactive analysis and assessment of single-cell copy-number variations. Nat. Methods 12:1058–1060.

Gawad C, Koh W, Quake SR. 2016. Single-cell genome sequencing: current state of the science. Nat. Rev. Genet. 17:175–188.

Goloboff PA, Catalano SA. 2016. TNT version 1.5, including a full implementation of phylogenetic morphometrics. Cladistics 32:221–238.

Huang L, Ma F, Chapman A, Lu S, Xie XS. 2015. Single-Cell Whole-Genome Amplification and Sequencing:Methodology and Applications. Annu. Rev. Genomics Hum. Genet. 16:79–102.

Jahn K, Kuipers J, Beerenwinkel N. 2016. Tree inference for single-cell data. Genome Biol. 17:86.

Kimura M. 1969. The number of heterozygous nucleotide sites maintained in a finite population due to steady flux of mutations. Genetics 61:893–903.

Kozlov AM, Darriba D, Flouri T, Morel B, Stamatakis A. 2019. RAxML-NG: A fast, scalable, and user-friendly tool for maximum likelihood phylogenetic inference. Bioinformatics 35:4453–4455.

Kozlov O. 2018. Models, optimizations, and tools for large-scale phylogenetic inference, handling sequence uncertainty, and taxonomic validation. Stamatakis A, Posada D, editors.

Kuipers J, Jahn K, Beerenwinkel N. 2017. Advances in understanding tumour evolution through single-cell sequencing. Biochim. Biophys. Acta Rev. Cancer 1867:127–138.

Kuipers J, Jahn K, Raphael BJ, Beerenwinkel N. 2017. Single-cell sequencing data reveal widespread recurrence and loss of mutational hits in the life histories of tumors. Genome Res. 27:1885–1894.

Lähnemann D, Köster J, Szczurek E, McCarthy DJ, Hicks SC, Robinson MD, Vallejos CA, Campbell KR, Beerenwinkel N, Mahfouz A, et al. 2020. Eleven grand challenges in single-cell data science. Genome Biol. 21:31.

Lee-Six H, Øbro NF, Shepherd MS, Grossmann S, Dawson K, Belmonte M, Osborne RJ, Huntly BJP, Martincorena I, Anderson E, et al. 2018. Population dynamics of normal human blood inferred from somatic mutations. Nature 561:473–478.

Lemoine F, Domelevo Entfellner J-B, Wilkinson E, Correia D, Dávila Felipe M, De Oliveira T, Gascuel O. 2018. Renewing Felsenstein’s phylogenetic bootstrap in the era of big data. Nature 556:452–456.

Leung ML, Davis A, Gao R, Casasent A, Wang Y, Sei E, Vilar E, Maru D, Kopetz S, Navin NE. 2017. Single-cell DNA sequencing reveals a late-dissemination model in metastatic colorectal cancer. Genome Res. 27:1287–1299.

Li H. 2011. A statistical framework for SNP calling, mutation discovery, association mapping and population genetical parameter estimation from sequencing data. Bioinformatics 27:2987–2993.

Li H. 2013. Aligning sequence reads, clone sequences and assembly contigs with BWA-MEM. arXiv [q-bio.GN] [Internet]. Available from: http://arxiv.org/abs/1303.3997

Li J, Zhang N, Zhang R, Sun L, Yu W, Guo W, Gao Y, Li M, Liu W, Liang P, et al. 2017. CDC5L Promotes hTERT Expression and Colorectal Tumor Growth. Cell. Physiol. Biochem. 41:2475–2488.

Lim B, Lin Y, Navin N. 2020. Advancing Cancer Research and Medicine with Single-Cell Genomics. Cancer Cell 37:456–470.

Li S-R, Gyselman VG, Lalude O, Dorudi S, Bustin SA. 2000. Transcription of the inositol polyphosphate 1-phosphatase gene (INPP1) is upregulated in human colorectal cancer. Molecular Carcinogenesis: Published in cooperation with the University of Texas MD Anderson Cancer Center 27:322–329.

Lutteropp S, Kozlov AM, Stamatakis A. 2020. A fast and memory-efficient implementation of the transfer bootstrap. Bioinformatics 36:2280–2281.

Macaulay IC, Voet T. 2014. Single cell genomics: advances and future perspectives. PLoS Genet. 10:e1004126.

Marioni JC, Arendt D. 2017. How Single-Cell Genomics Is Changing Evolutionary and Developmental Biology. Annu.Rev. Cell Dev. Biol. 33:537–553.

Martincorena I, Campbell PJ. 2015. Somatic mutation in cancer and normal cells. Science 349:1483–1489.

Ma SSQ, Srivastava S, Llamosas E, Hawkins NJ, Hesson LB, Ward RL, Ford CE. 2016. ROR2 is epigenetically inactivated in the early stages of colorectal neoplasia and is associated with proliferation and migration. BMC Cancer 16:508.

Navin NE. 2014. Cancer genomics: one cell at a time. Genome Biol. 15:452.

Pan H, Pan J, Song S, Ji L, Lv H, Yang Z. 2019. EXOSC5 as a Novel Prognostic Marker Promotes Proliferation of Colorectal Cancer via Activating the ERK and AKT Pathways. Front. Oncol. 9:643.

Paradis E, Schliep K. 2019. ape 5.0: an environment for modern phylogenetics and evolutionary analyses in R. Bioinformatics 35:526–528.

Poplin R, Ruano-Rubio V, DePristo MA, Fennell TJ. 2018. Scaling accurate genetic variant discovery to tens of thousands of samples. BioRxiv [Internet]. Available from: https://www.biorxiv.org/content/10.1101/201178v3.abstract

Posada D. 2020. CellCoal: Coalescent Simulation of Single-Cell Sequencing Samples. Mol. Biol. Evol. 37:1535–1542.

Robinson DF. 1971. Comparison of labeled trees with valency three. J. Combin. Theory Ser. B 11:105–119.

Robinson DF, Foulds LR. 1981. Comparison of phylogenetic trees. Math. Biosci. 53:131–147.

Romero-Pérez L, Surdez D, Brunet E, Delattre O, Grünewald TGP. 2019. STAG Mutations in Cancer. Trends Cancer Res. 5:506–520.

Ross EM, Markowetz F. 2016. OncoNEM: inferring tumor evolution from single-cell sequencing data. Genome Biol. 17:69.

Satas G, Zaccaria S, Mon G, Raphael BJ. 2020. SCARLET: Single-Cell Tumor Phylogeny Inference with Copy-Number Constrained Mutation Losses. Cell Systems 10:323–332.e8.

Singer J, Kuipers J, Jahn K, Beerenwinkel N. 2018. Single-cell mutation identification via phylogenetic inference. Nat. Commun. 9:5144.

Stamatakis A. 2014. RAxML version 8: a tool for phylogenetic analysis and post-analysis of large phylogenies. Bioinformatics 30:1312–1313.

Stegle O, Teichmann SA, Marioni JC. 2015. Computational and analytical challenges in single-cell transcriptomics. Nat. Rev. Genet. 16:133–145.

Tanay A, Regev A. 2017. Scaling single-cell genomics from phenomenology to mechanism. Nature 541:331.

Tavaré S. 1986. Some probabilistic and statistical problems in the analysis of DNA sequences. Lectures on mathematics in the life sciences 17:57–86.

Wickham H. 2016. ggplot2: Elegant Graphics for Data Analysis. Springer

Wiedmeier JE, Noel P, Lin W, Von Hoff DD, Han H. 2019. Single-Cell Sequencing in Precision Medicine. Precision Medicine in Cancer Therapy: 237–252.

Yang Z, Kumar S, Nei M. 1995. A new method of inference of ancestral nucleotide and amino acid sequences. Genetics 141:1641–1650.

Yu G, Smith DK, Zhu H, Guan Y, Lam TT. 2017. ggtree : an r package for visualization and annotation of phylogenetic trees with their covariates and other associated data. Methods in Ecology and Evolution 8:28–36.

Zafar H, Navin N, Chen K, Nakhleh L. 2019. SiCloneFit: Bayesian inference of population structure, genotype, and phylogeny of tumor clones from single-cell genome sequencing data. Genome Res. 29:1847–1859.

Zafar H, Navin N, Nakhleh L, Chen K. 2018. Computational approaches for inferring tumor evolution from single-cell genomic data. Current Opinion in Systems Biology 7:16–25.

Zafar H, Tzen A, Navin N, Chen K, Nakhleh L. 2017. SiFit: inferring tumor trees from single-cell sequencing data under finite-sites models. Genome Biol. 18:178.

Zafar H, Wang Y, Nakhleh L, Navin N, Chen K. 2016. Monovar: single-nucleotide variant detection in single cells. Nat. Methods 13:505–507.

Zhou X, Shen X-X, Hittinger CT, Rokas A. 2018. Evaluating Fast Maximum Likelihood-Based Phylogenetic Programs Using Empirical Phylogenomic Data Sets. Mol. Biol. Evol. 35:486–503.

