## Supplementary Material for "CellPhy: accurate and fast probabilistic inference of single-cell phylogenies from scDNA-seq data"

**Contents**

Table S1. Simulation scenarios.

Table S2. Genotype codes.

Table S3. CellPhy's compatibility with variant callers.

Figure S1. Phylogenetic accuracy in Simulation 1 ("target-ISM") with 500 SNVs.

Figure S2. Phylogenetic accuracy in Simulation 1 ("target-ISM") with 1000 SNVs.

Figure S3. Phylogenetic reconstruction accuracy in Simulation 3 ("WGS-sig") with signature S1.

Figure S4. Phylogenetic reconstruction accuracy in Simulation 3 ("WGS-sig") with signature S5.

Figure S5. Phylogenetic accuracy in Simulation 4 ("NGS-like") at 30x.

Figure S6. Phylogenetic accuracy in Simulation 4 ("NGS-like") at 100x.

Figure S7. Phylogenetic accuracy in Simulation 5 ("NGS-doublet").

Figure S8. Estimation of the genotype error and ADO rate.

Figure S9. SiFIT, SCIPhI, infSCITE, and TNT trees for the CRC24 dataset.

Figure S10. Non-synonymous mutations along the CellPhy L86 tree.

Figure S11. SiFIT, SCIPhI, infSCITE, and TNT trees for the L86 dataset.

Figure S12. Phylogenetic reconstruction of 15 whole-genome sequenced neurons from a healthy donor.

Figure S13. Phylogenetic reconstruction of 140 whole-genome sequenced single-cell derived hematopoietic "colonies" from a healthy donor.

Figure S14. Genotype likelihood vectors for a simple unrooted phylogenetic tree.

Figure S15. Phylogenetic likelihood calculations.

Figure S16. Phylogenetic bootstrap calculation.

Figure S17. Phylogenetic accuracy with an approximate model in Simulation 6 ("NGS-large").

Supplementary Note 1. Genotype error model

Supplementary Note 2. Approximate model of evolution for unphased diploid genotypes

Supplementary Note 3. Standard phylogenetic likelihood calculations on DNA sequence alignments

**Table S1. Simulation scenarios.**

| | **Sim 1 target-ISM** | **Sim 2 WGS-FSM** | **Sim 3 WGS-sig** | **Sim 4 NGS-like** | **Sim 5 NGS-doublet** | **Sim 6 NGS-large** |
|---|---|---|---|---|---|---|
| Number of cells | 40 | 100 | 60 | 40 | 40 | 100, 500, 1000 |
| Number of sites | 5000 | 10000 | 10000 | 10000 | 10000 | 1000, 10000, 50000 |
| Effective population size | 10000 | 10000 | 10000 | 10000 | 10000 | 10000 |
| Exponential growth rate | $10^{-4}$ | $10^{-4}$ | $10^{-4}$ | $10^{-4}$ | $10^{-4}$ | $10^{-4}$ |
| Root branch length | 0.01 | 0.01 | 0.01 | 0.01 | 0.01 | 0.01 |
| Outgroup branch length | 0 | 0 | 0 | 0 | 0 | 0 |
| Lineage rate variation (alpha) | 1.0 | 1.0 | 1.0 | 1.0 | 1.0 | 1.0 |
| Number of fixed SNVs | 250, 500, 1000 | n/a | n/a | n/a | n/a | n/a |
| Number of true SNVs | 250, 500, 1000 | 1695 - 2560 | 1414 - 4215 | 1376 - 2123 | 1272 - 2094 | 168-335, XXX, XXX |
| Number of observed SNVs | 206-4995, 415-4956, 807-4961 | 1531 - 10000 | 1224 - 10000 | 1147 - 10000, 1226 - 6841, 1227 - 2828 | 1271 - 9963 | 1000 / 10000 / 50000 |
| Mutation rate | n/a | $10^{-6}$ | $10^{-6}$ | $10^{-6}$ | $10^{-6}$ | $10^{-6}$ |
| Mutation model | ISM diploid | GTnR | ISM S1 and S5 | GTnR | GTnR | GTnR |
| Mutation rate variation among sites (alpha) | n/a | 1.0 | n/a | 1.0 | 1.0 | 1.0 |
| Genotype error | 0, 0.01, 0.05, 0.10 | 0, 0.01, 0.05, 0.10 | 0, 0.01, 0.10, 0.20 | n/a | n/a | n/a |

| | | | | | | |
|---|---|---|---|---|---|---|
| ADO | 0, 0.10, 0.25, 0.50 | 0, 0.10, 0.25, 0.50 | 0, 0.05, 0.15, 0.50 | 0, 0.10, 0.25 | 0, 0.10 | 0.10 |
| Sequencing depth | n/a | n/a | n/a | 5, 30, 100 | 5 | 5 |
| Coverage overdispersion | n/a | n/a | n/a | 5 | 5 | 5 |
| Sequencing error | n/a | n/a | n/a | 0, 0.01, 0.05 | 0/0.01 | 0.01 |
| Amplification error mean* | n/a | n/a | n/a | 0, 0.05, 0.10 | 0/0.05 | 0.05 |
| Amplification error variance | n/a | n/a | n/a | 0.01 | 0.01 | 0.01 |
| Allelic imbalance | n/a | n/a | n/a | 0.50 | 0.50 | 0.50 |
| Haploid coverage reduction | n/a | n/a | n/a | 0.50 | 0.50 | 0.50 |
| Doublet rate | 0 | 0 | 0 | 0 | 0, 0.05, 0.10, 0.20 | 0 |
| Number of replicates | 100 | 100 | 100 | 100 | 100 | 20 |

**Table S2. Genotype codes.** Encoding of unphased genotypes as single letters for the FASTA/PHYLIP genotype files and translation to phased genotype likelihoods.

<table>
<tr><th colspan="2">Symbol</th><th>A</th><th>C</th><th>G</th><th>T</th><th>M</th><th>R</th><th>W</th><th>S</th><th>Y</th><th>K</th></tr>
<tr><td colspan="2">Genotype (unphased )</td><td>A/A</td><td>C/C</td><td>G/G</td><td>T/T</td><td>A/C</td><td>A/G</td><td>A/T</td><td>C/G</td><td>C/T</td><td>G/T</td></tr>
<tr><td colspan="2">Genotypes (phased)</td><td>A|A</td><td>C|C</td><td>G|G</td><td>T|T</td><td>A|C<br>C|A</td><td>A|G<br>G|A</td><td>A|T<br>T|A</td><td>C|G<br>G|C</td><td>C|T<br>T|C</td><td>G|T<br>T|G</td></tr>
<tr><td rowspan="16">Phased genotype likelihood</td><td>A|A</td><td>1</td><td>0</td><td>0</td><td>0</td><td>0</td><td>0</td><td>0</td><td>0</td><td>0</td><td>0</td></tr>
<tr><td>C|C</td><td>0</td><td>1</td><td>0</td><td>0</td><td>0</td><td>0</td><td>0</td><td>0</td><td>0</td><td>0</td></tr>
<tr><td>G|G</td><td>0</td><td>0</td><td>1</td><td>0</td><td>0</td><td>0</td><td>0</td><td>0</td><td>0</td><td>0</td></tr>
<tr><td>T|T</td><td>0</td><td>0</td><td>0</td><td>1</td><td>0</td><td>0</td><td>0</td><td>0</td><td>0</td><td>0</td></tr>
<tr><td>A|C</td><td>0</td><td>0</td><td>0</td><td>0</td><td>1</td><td>0</td><td>0</td><td>0</td><td>0</td><td>0</td></tr>
<tr><td>A|G</td><td>0</td><td>0</td><td>0</td><td>0</td><td>0</td><td>1</td><td>0</td><td>0</td><td>0</td><td>0</td></tr>
<tr><td>A|T</td><td>0</td><td>0</td><td>0</td><td>0</td><td>0</td><td>0</td><td>1</td><td>0</td><td>0</td><td>0</td></tr>
<tr><td>C|G</td><td>0</td><td>0</td><td>0</td><td>0</td><td>0</td><td>0</td><td>0</td><td>1</td><td>0</td><td>0</td></tr>
<tr><td>C|T</td><td>0</td><td>0</td><td>0</td><td>0</td><td>0</td><td>0</td><td>0</td><td>0</td><td>1</td><td>0</td></tr>
<tr><td>G|T</td><td>0</td><td>0</td><td>0</td><td>0</td><td>0</td><td>0</td><td>0</td><td>0</td><td>0</td><td>1</td></tr>
<tr><td>C|A</td><td>0</td><td>0</td><td>0</td><td>0</td><td>1</td><td>0</td><td>0</td><td>0</td><td>0</td><td>0</td></tr>
<tr><td>G|A</td><td>0</td><td>0</td><td>0</td><td>0</td><td>0</td><td>1</td><td>0</td><td>0</td><td>0</td><td>0</td></tr>
<tr><td>T|A</td><td>0</td><td>0</td><td>0</td><td>0</td><td>0</td><td>0</td><td>1</td><td>0</td><td>0</td><td>0</td></tr>
<tr><td>G|C</td><td>0</td><td>0</td><td>0</td><td>0</td><td>0</td><td>0</td><td>0</td><td>1</td><td>0</td><td>0</td></tr>
<tr><td>T|C</td><td>0</td><td>0</td><td>0</td><td>0</td><td>0</td><td>0</td><td>0</td><td>0</td><td>1</td><td>0</td></tr>
<tr><td>T|G</td><td>0</td><td>0</td><td>0</td><td>0</td><td>0</td><td>0</td><td>0</td><td>0</td><td>0</td><td>1</td></tr>
</table>

**Table S3. CellPhy's compatibility with variant callers.**

| Algorithm | Single-cell specific | Multi-sample calling | Output format | CellPhy support | | Reference |
|---|---|---|---|---|---|---|
| | | | | ML mode | GL mode | |
| SC-Caller | Yes | No | (VCF) | Yes | Script | Dong et al. 2017 |
| Monovar | Yes | Yes | VCF | Yes | Yes | Zafar et al. 2016 |
| Prosolo | Yes | No | BCF | Yes | Yes | Lähnemann et al. 2020 |
| Conbase | Yes | Yes | TSV | (Script) | No | Hård et al. 2019 |
| SCIPhI | Yes | Yes | (VCF) | Yes | No | Singer et al. 2018 |
| scan-snv | Yes | No | RDA | (Script) | No | Luquette et al. 2019 |
| HaplotypeCaller | No | Yes | VCF | Yes | Yes | Poplin et al. 2018 |

Remarks:

| | |
|---|---|
| | Although not single-cell specific, HaplotypeCaller is still widely popular within the single-cell genomics community. |
| (VCF) | Non-standard VCF |
| Script | Conversion script provided with CellPhy |
| (Script) | Conversion possible with a custom script (not provided) |

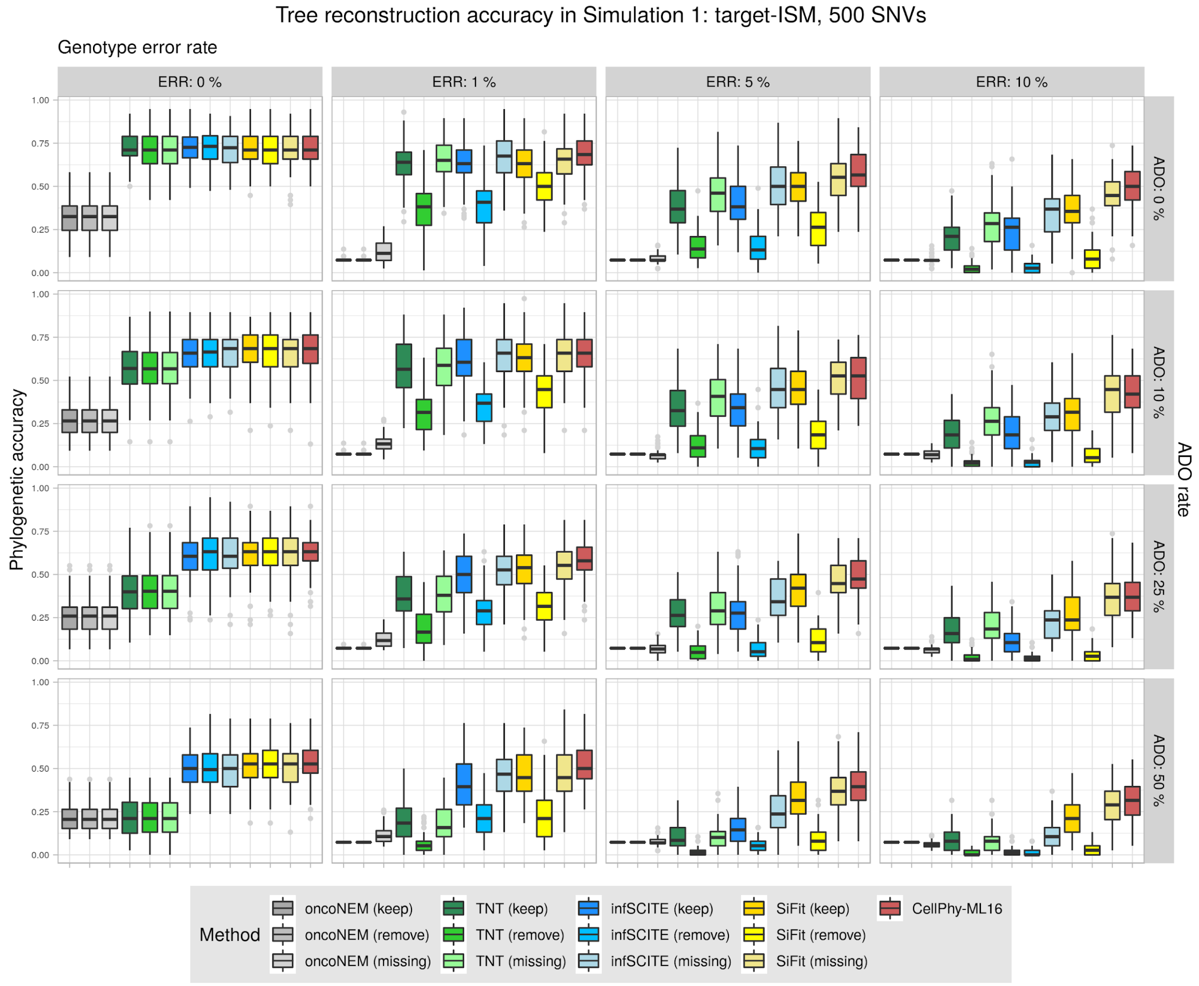


**Figure S1. Phylogenetic accuracy in Simulation 1 ("target-ISM") with 500 SNVs**. Datasets consisted of 40 cells. Accuracy was evaluated under different levels of genotype error (ERR), allelic dropout (ADO), and genotype recoding strategies ("keep", "remove", "missing") as explained in the main text. Phylogenetic accuracy is defined as 1 – nRF (see Methods). Boxplots were generated using the ggplot2 R package (https://ggplot2.tidyverse.org) with default parameters. The lower and upper hinges correspond to the first and third quartiles. The upper whisker extends from the hinge to the largest value no further than 1.5 * IQR from the hinge (where IQR is the interquartile range or distance between the first and third quartiles). The lower whisker extends from the hinge to the smallest value at most 1.5 * IQR of the hinge. Data beyond the end of the whiskers are called "outlying" points and are plotted individually.

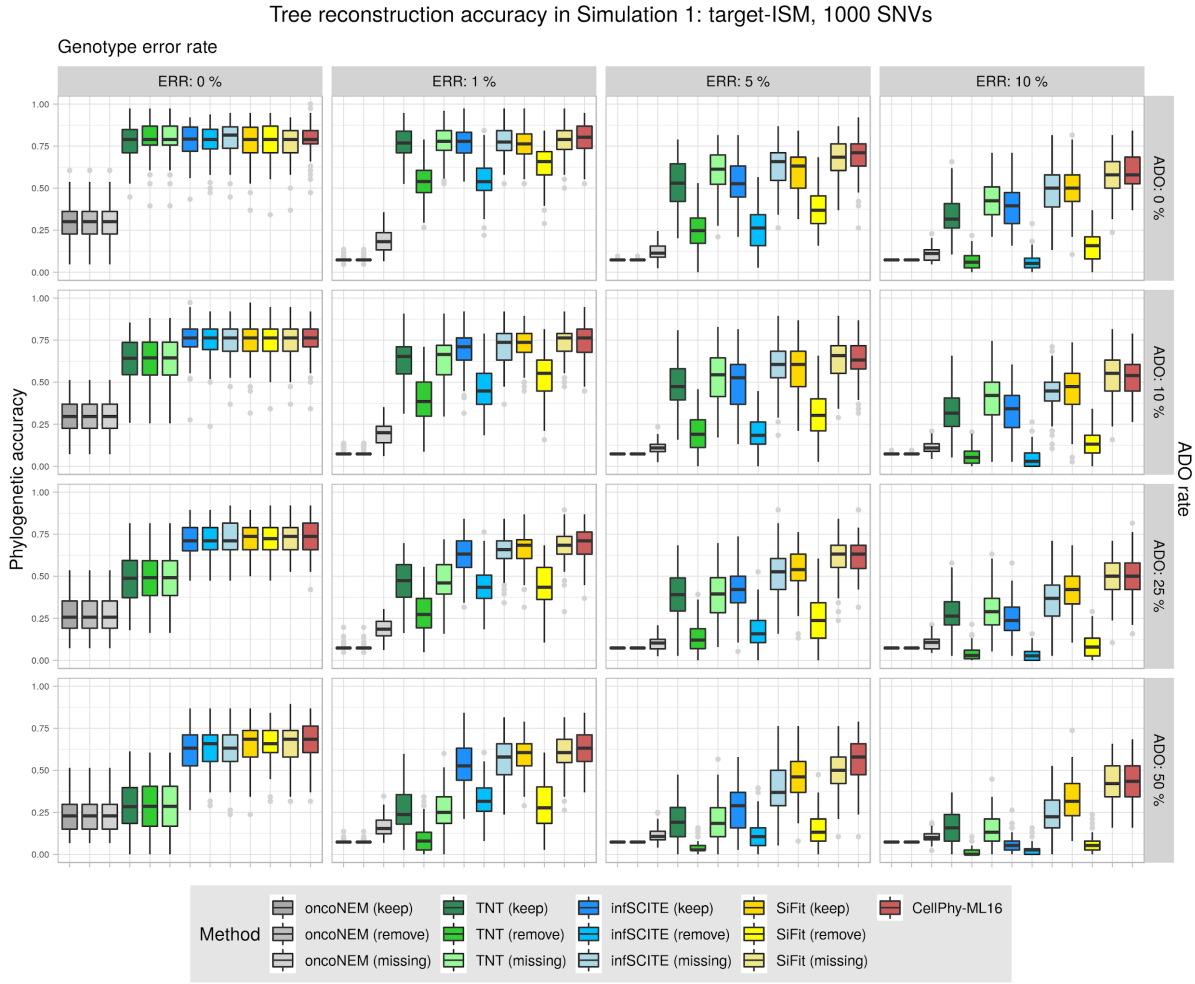


**Figure S2. Phylogenetic accuracy in Simulation 1 (“target-ISM”) with 1000 SNVs.** Datasets consisted of 40 cells. Accuracy was evaluated under different levels of genotype error (ERR), allelic dropout (ADO), and genotype recoding strategies (“keep”, “remove”, “missing”) as explained in the main text. Phylogenetic accuracy is defined as 1 – nRF (see Methods). See Figure S1 for an explanation of the boxplots.

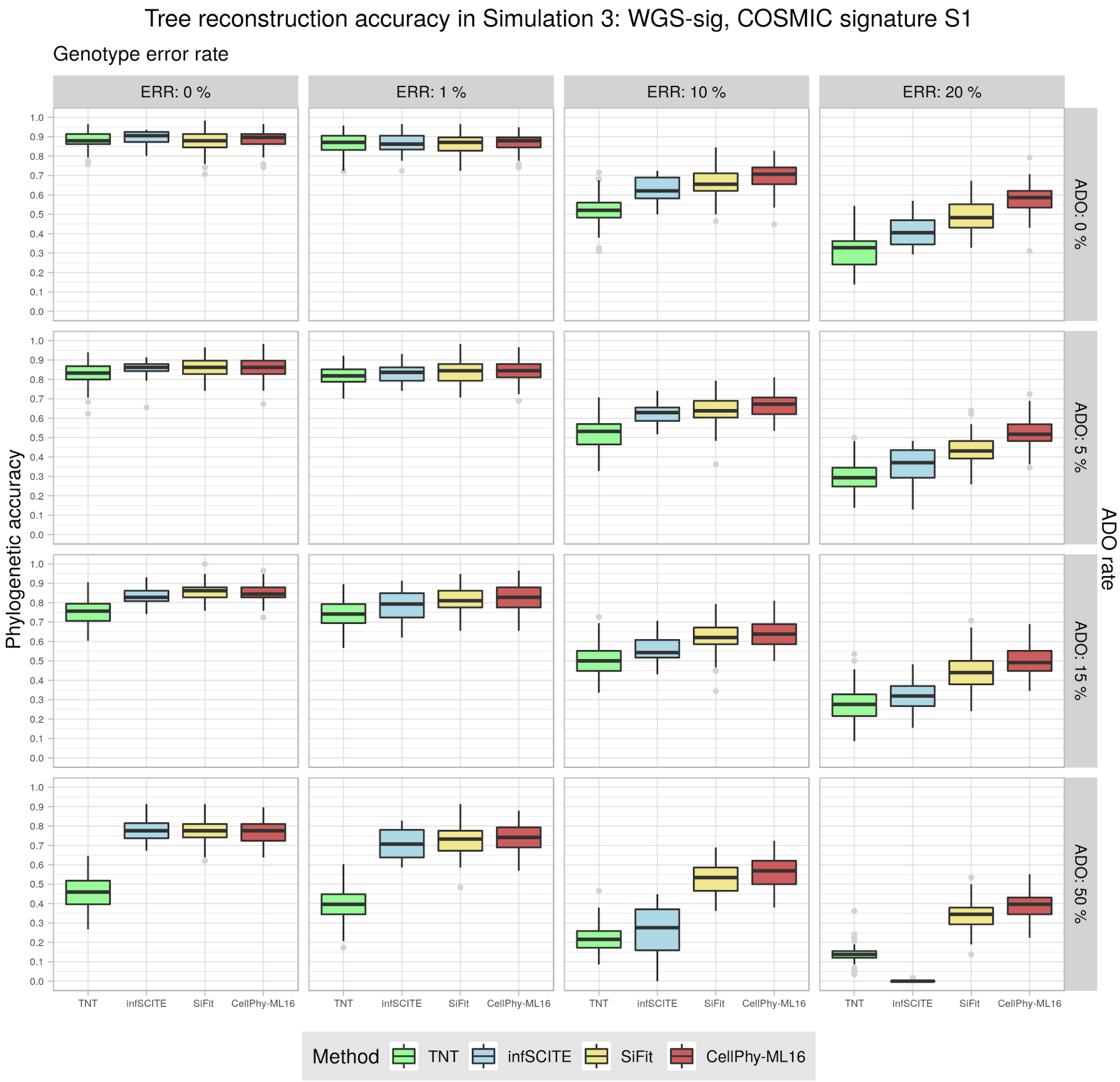


**Figure S3. Phylogenetic reconstruction accuracy in Simulation 3 ("WGS-sig") with signature S1.** Datasets consisted of 60 cells and 1000-4000 SNVs. Accuracy was evaluated under different levels of genotype error (ERR) and allelic dropout (ADO). Phylogenetic accuracy is defined as 1 – nRF (see Methods). See Figure S1 for an explanation of the boxplots.

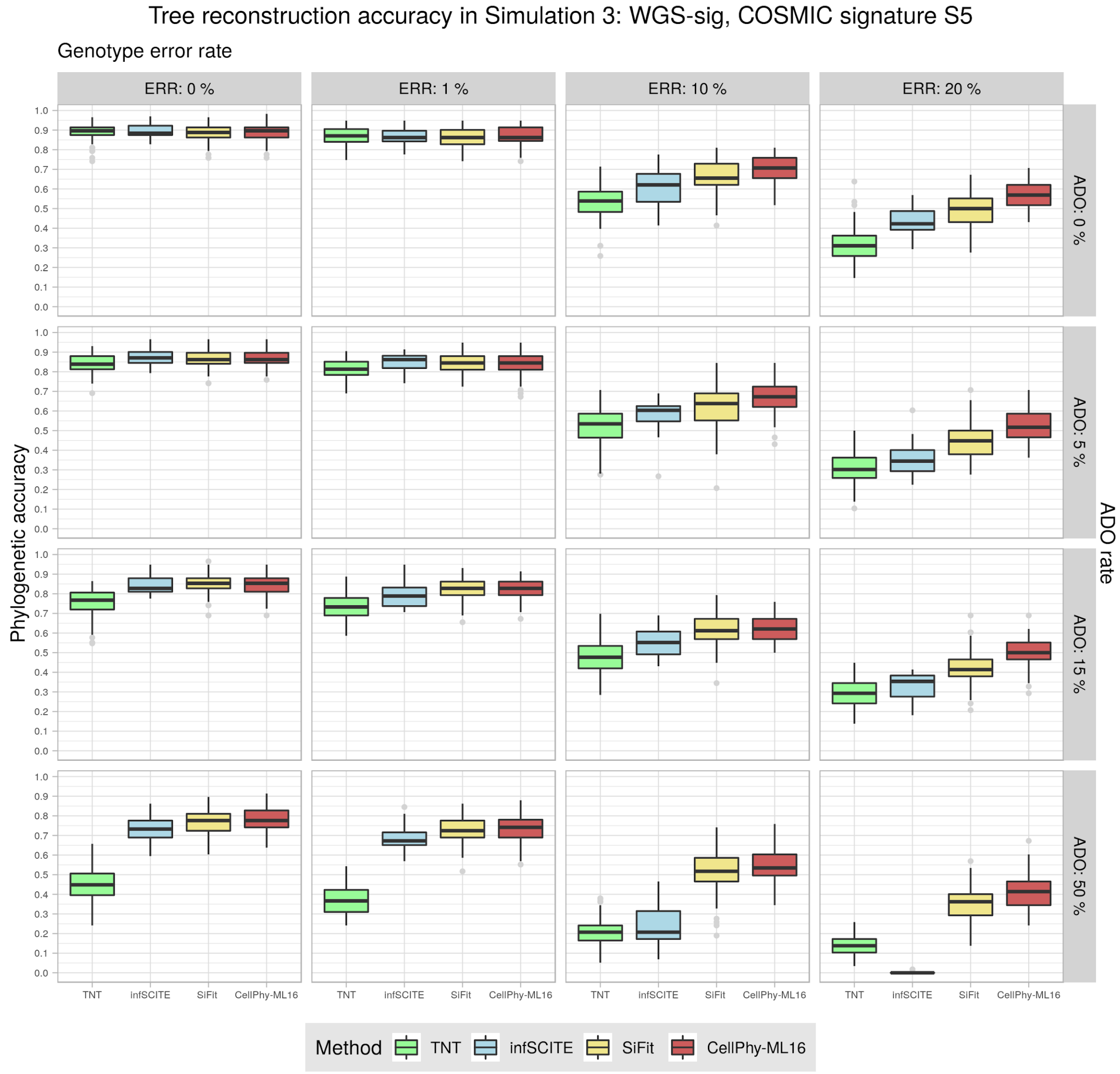


**Figure S4. Phylogenetic reconstruction accuracy in Simulation 3 ("WGS-sig") with signature S5.** Datasets consisted of 60 cells and ~1000-4000 SNVs. Accuracy was evaluated under different levels of genotype error (ERR) and allelic dropout (ADO). Phylogenetic accuracy is defined as 1 – nRF (see Methods). See Figure S1 for an explanation of the boxplots.

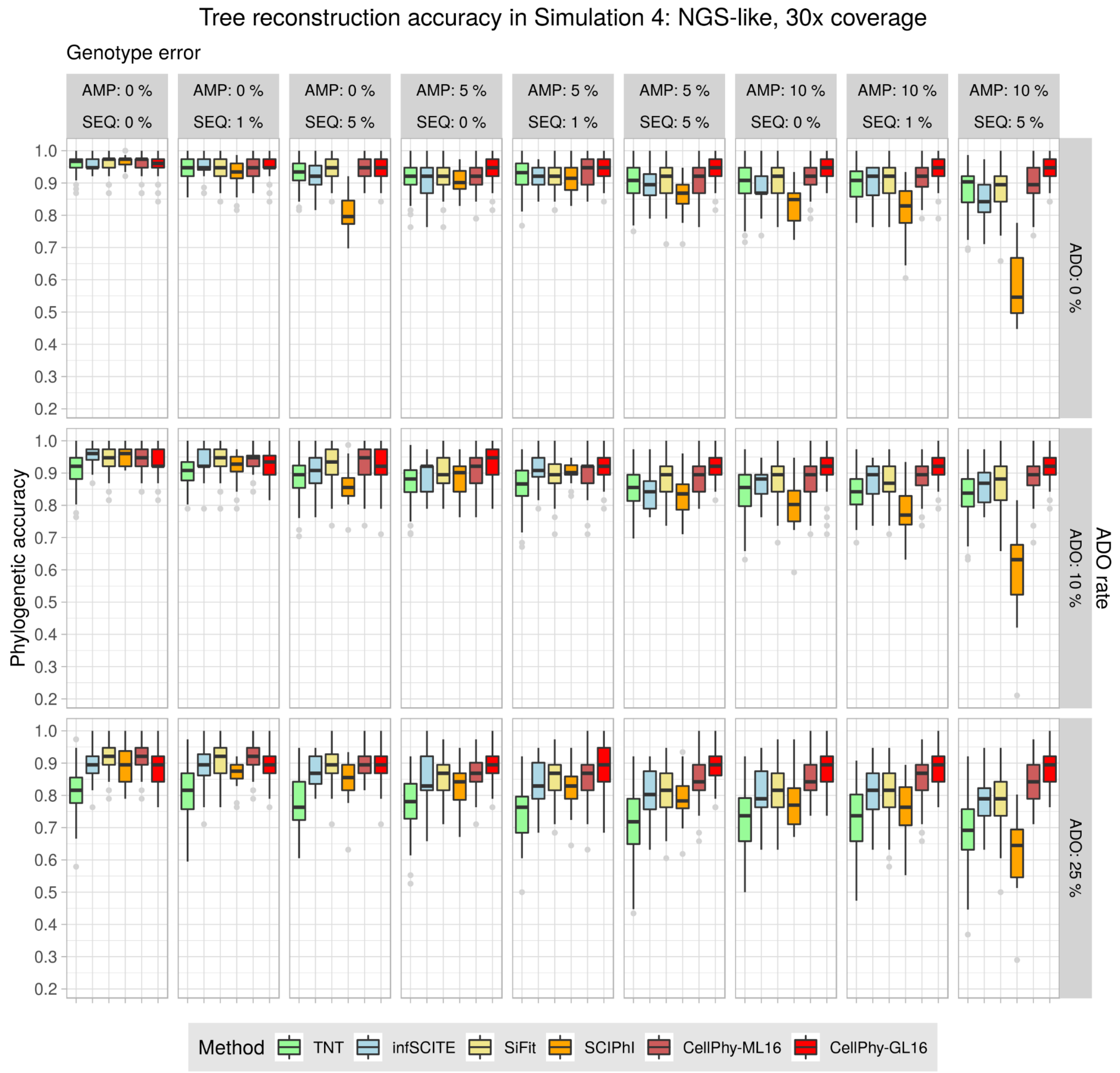


**Figure S5**. **Phylogenetic accuracy in Simulation 4 ("NGS-like") at 30x**. Datasets consisted of 40 cells and ~1000-2000 SNVs. All methods use the ML genotypes except CellPhy-GL, which uses the genotype likelihoods. Phylogenetic accuracy is defined as 1 – nRF (see Methods). AMP is the amplification error rate, SEQ is the sequencing error rate, and ADO is the allelic dropout rate. See Figure S1 for an explanation of the boxplots.

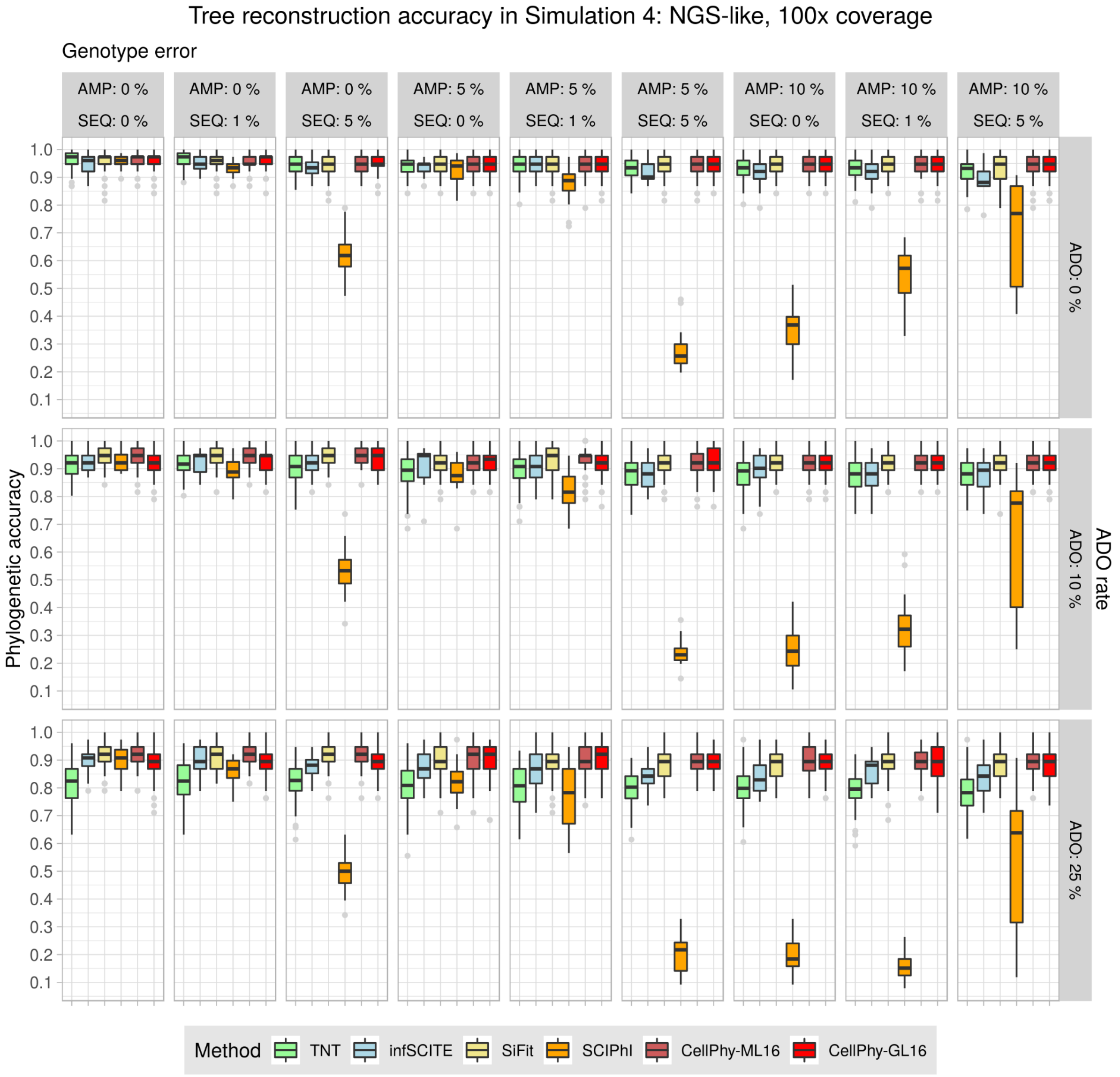


**Figure S6. Phylogenetic accuracy in Simulation 4 ("NGS-like") at 100x**. Datasets consisted of 40 cells and ~1000-2000 SNVs. All methods use the ML genotypes except CellPhy-GL, which uses the genotype likelihoods. Phylogenetic accuracy is defined as 1 – nRF (see Methods). AMP is the amplification error rate, SEQ is the sequencing error rate, and ADO is the allelic dropout rate. See Figure S1 for an explanation of the boxplots.

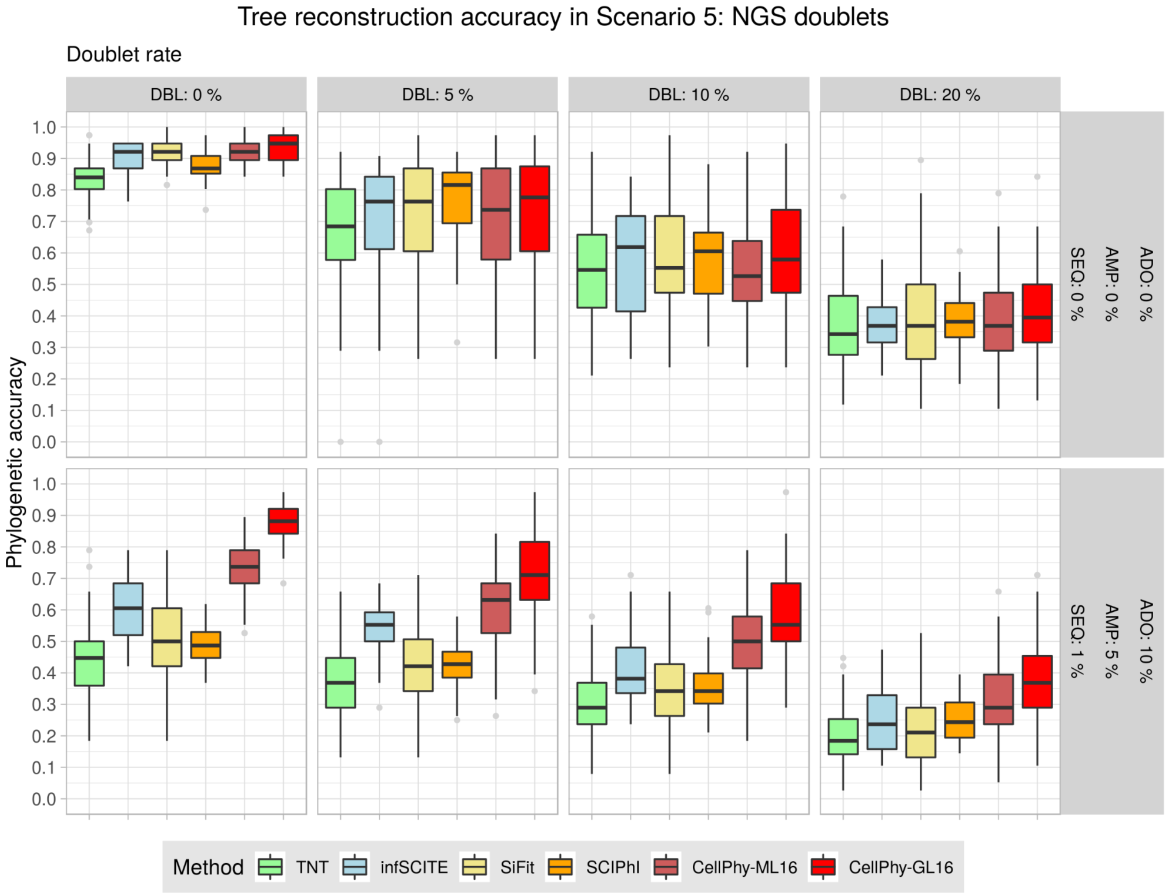


**Figure S7. Phylogenetic accuracy in Simulation 5 ("NGS-doublet")**. Data simulated under mutational signature S1 and with a 5x sequencing depth. All methods use the ML genotypes except CellPhy-GL, which uses the genotype likelihoods. Phylogenetic accuracy is defined as 1 – nRF (see Methods). DBL is the double rate, AMP is the amplification error rate, SEQ is the sequencing error rate, and ADO is the allelic dropout rate. See Figure S1 for an explanation of the boxplots.

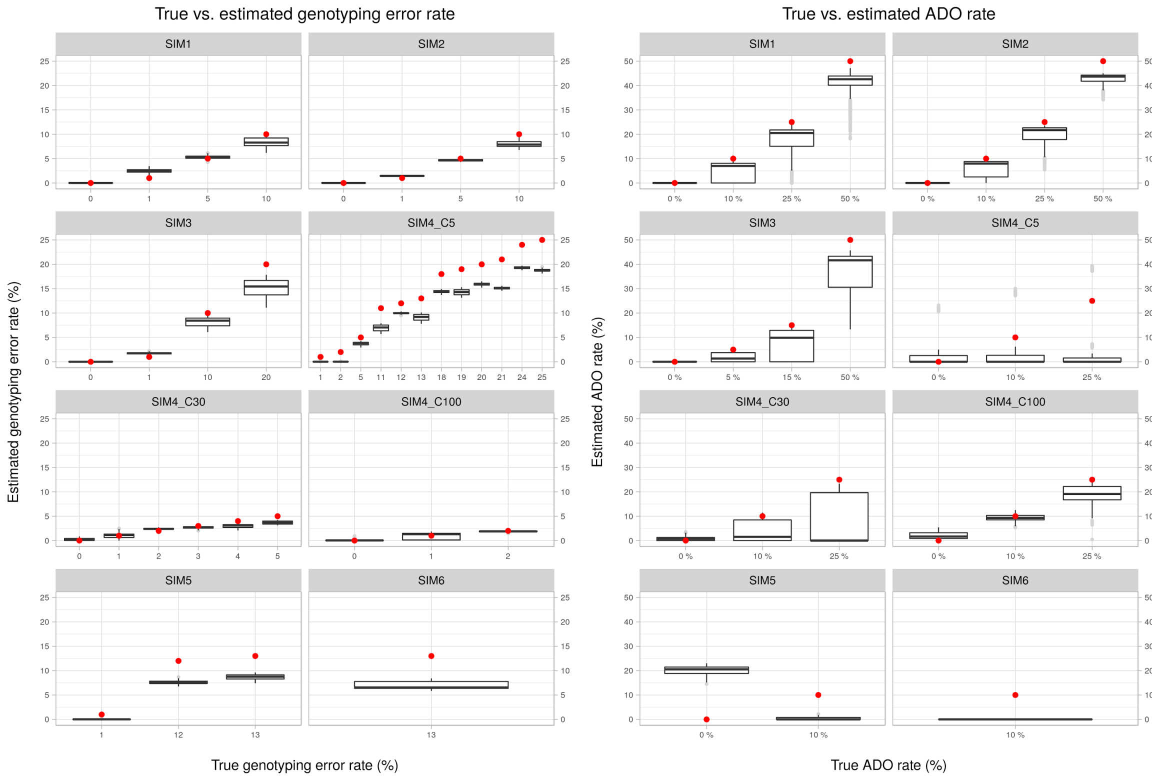


**Figure S8. Estimation of the genotype error and ADO rate.** (A) Genotype error ML estimates. (B) ADO rate ML estimates. For simulation 1, 250, 500 or 1,000 SNVs were considered. Red dots highlight the true values. See Figure S1 for an explanation of the boxplots.

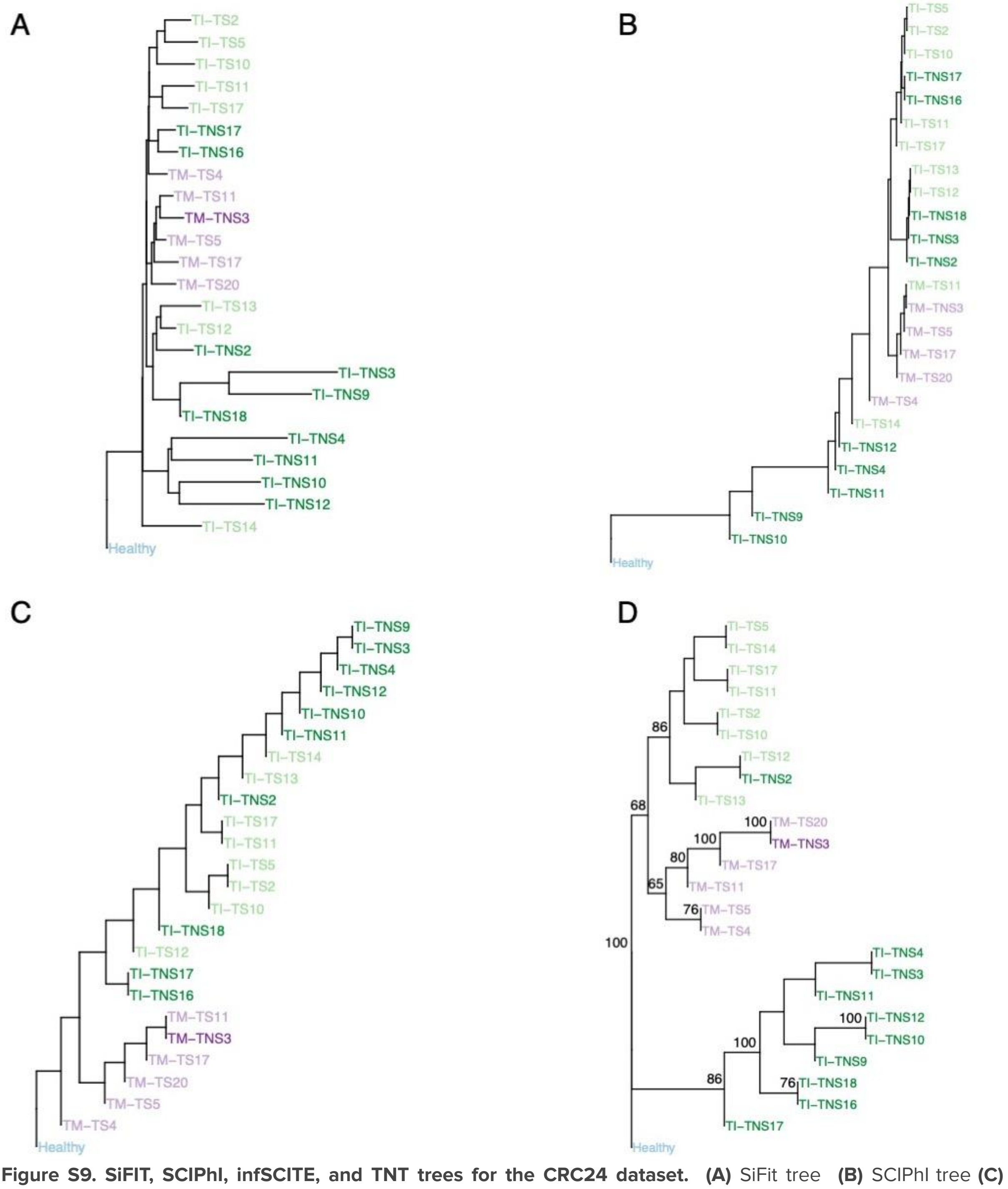


**Figure S9. SiFIT, SCIPhI, infSCITE, and TNT trees for the CRC24 dataset.** **(A)** SiFit tree **(B)** SCIPhI tree **(C)** infSCITE tree **(D)** TNT tree**.** Distinct colors represent cell type: healthy (blue); tumor-non-stem from TI region (dark green), tumor-stem from TI region (light green), tumor-non-stem from TM region (dark purple), tumor-stem from TM region (light purple). Only bootstrap support values above 50 are shown.

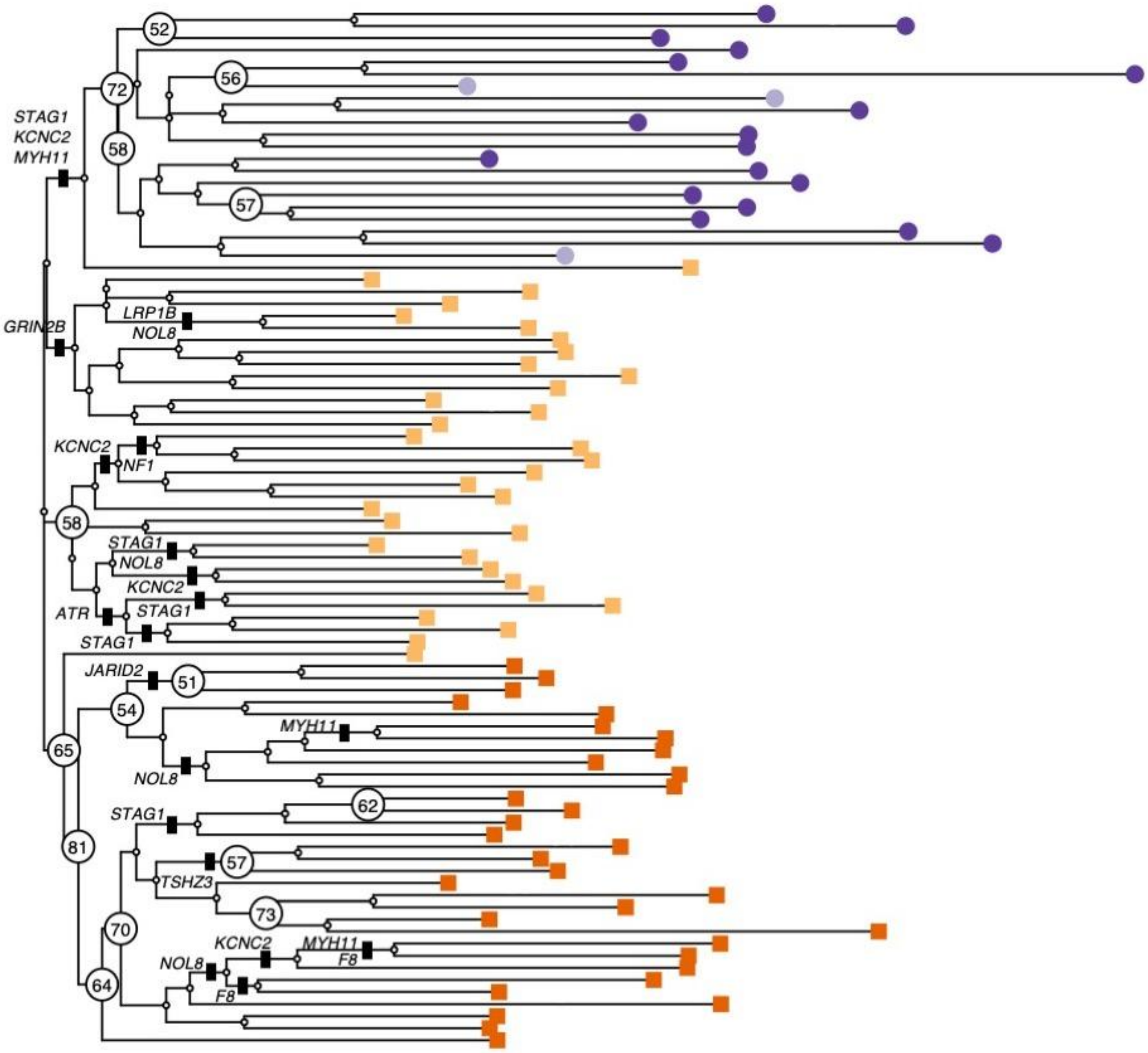


**Figure S10. Non-synonymous mutations along the CellPhy L86 tree.** The figure shows the CellPhy L86 tree with non-synonymous mutations mapped to the internal branches of the tree. Distinct shapes and colors represent cell type: healthy diploid cells - from both primary and metastatic sites - (dark purple circle), healthy diploid cells missorted (light purple circle), primary tumor aneuploid cells (light orange square), metastatic aneuploid cells (dark orange square). Only bootstrap values above 50 are shown.

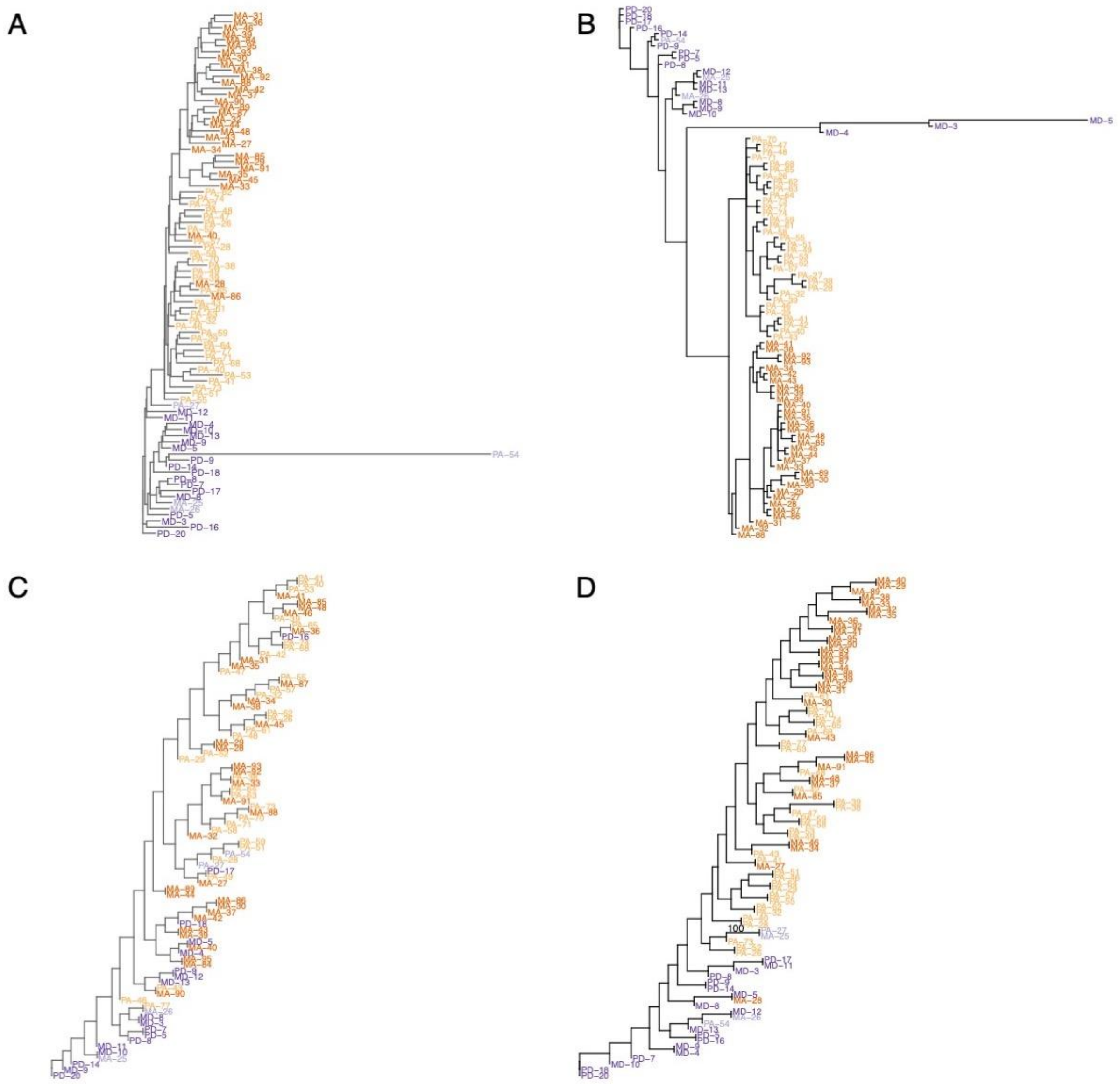


**Figure S11. SiFIT, SCIPhI, infSCITE, and TNT trees for the L86 dataset. (A)** SiFit tree **(B)** SCIPhI tree **(C)** infSCITE tree **(D)** TNT tree. Distinct colors represent cell type: healthy diploid cells from both primary and metastatic sites (dark purple), healthy diploid cells missorted (light purple), primary tumor aneuploid cells (light orange), metastatic aneuploid cells (dark orange). Only bootstrap values above 50 are shown.

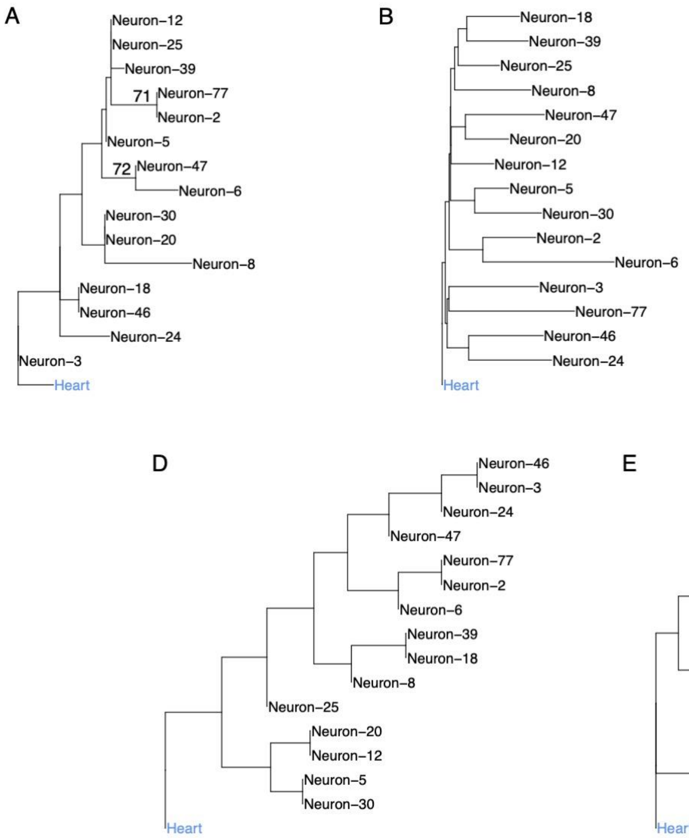


**Figure S12. Phylogenetic reconstruction of 15 whole-genome sequenced neurons from a healthy donor. (A)** Single-cell ML phylogenetic tree inferred with "Cellphy-GL". Only bootstrap support values above 50 are shown. **(B)** SiFit ML phylogenetic tree using a ternary genotype matrix. **(C)** SCIPhI ML phylogenetic tree. **(D)** infSCITE ML phylogenetic tree using a binary genotype matrix. **(E)** TNT MP phylogenetic tree using binary alignments. Only bootstrap values above 50 are shown.

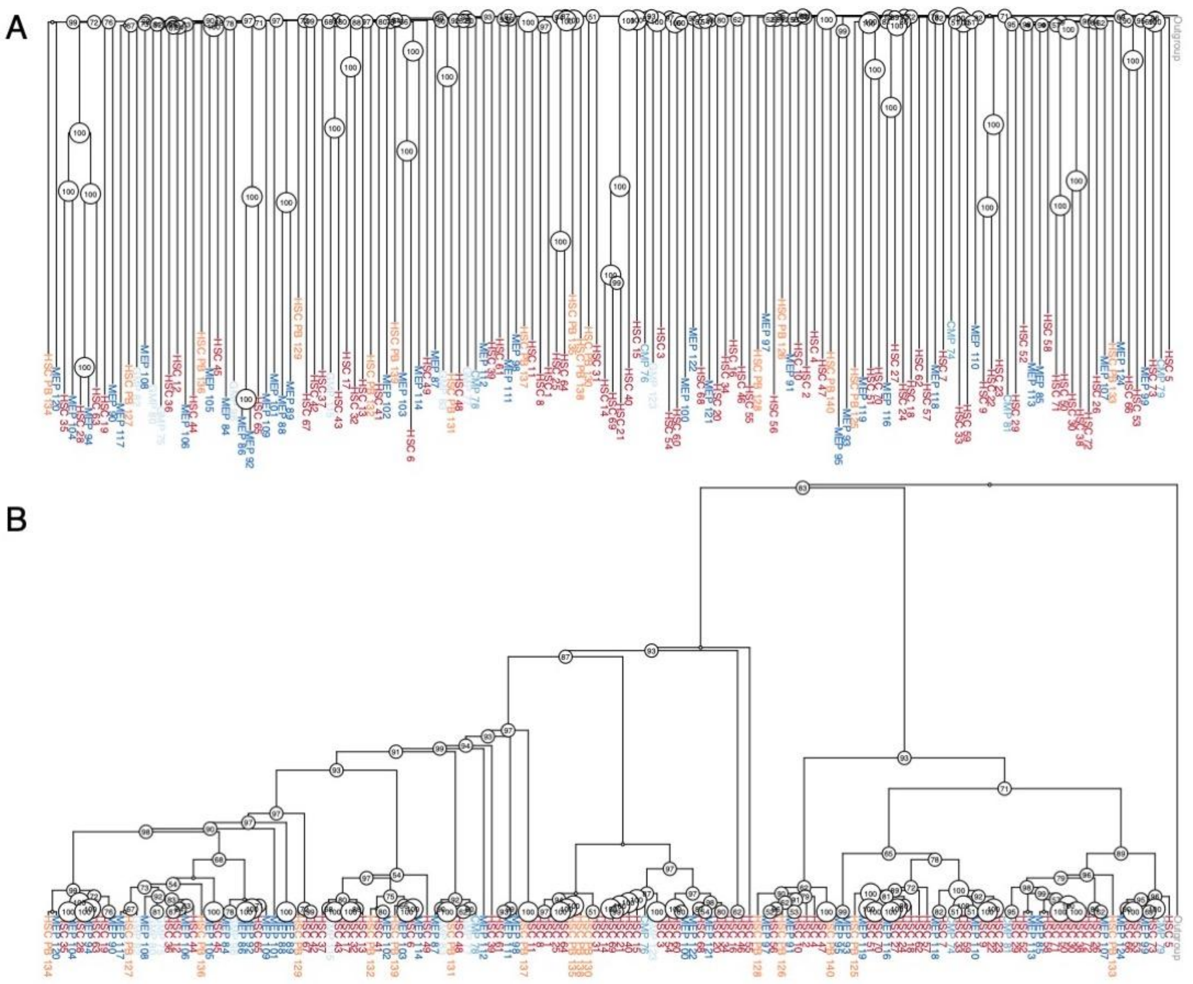

**Figure S13. Phylogenetic reconstruction of 140 whole-genome sequenced single-cell derived hematopoietic "colonies" from a healthy donor.** (A) ML phylogenetic tree inferred with Cellphy using a genotype matrix with 127,884 mutations. Only bootstrap support values above 50 are shown. (B) The same ML phylogeny as in (A), but ignoring branch lengths to ease visualization of ancestral relationships. Only bootstrap values above 50 are shown.

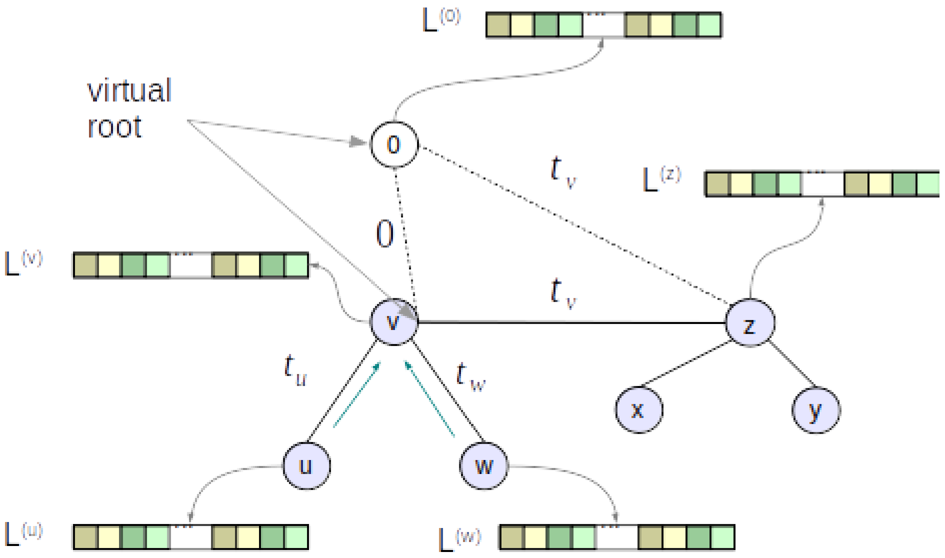


**Figure S14**. Genotype likelihood vectors for a simple unrooted phylogenetic tree. To compute the likelihood of the tree via the Felsenstein pruning algorithm, we place a virtual root (*node 0*) at an arbitrary inner node (here: *node v*). Then, we perform a post-order tree traversal and compute likelihood vectors at each inner node recursively according to equation (12). We use vectors $L^u$ and $L^w$to compute $L^v$, vectors $L^x$ and $L^y$ (not shown) to compute $L^z$, and then finally vectors $L^v$ and $L^z$to compute $L^0$.

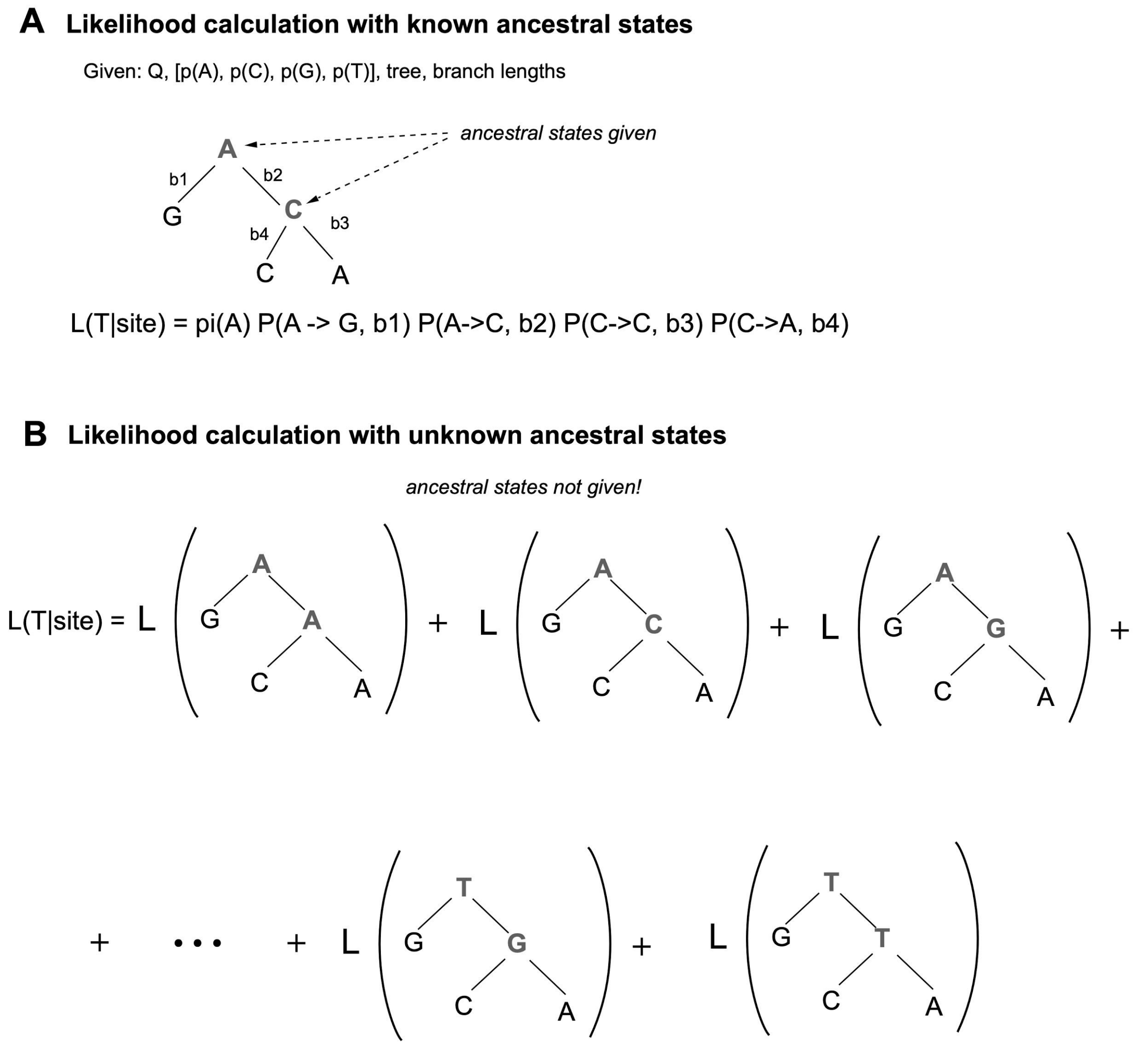


**Figure S15. Phylogenetic likelihood calculations. (A)** Outline of a simple likelihood calculation on a given tree with given branch lengths, when the inner states are also given. **(B)** The likelihood is just the sum over the likelihoods of *all possible* evolutionary scenarios. The likelihood of the trees in parentheses is calculated as in the simple example for given states.

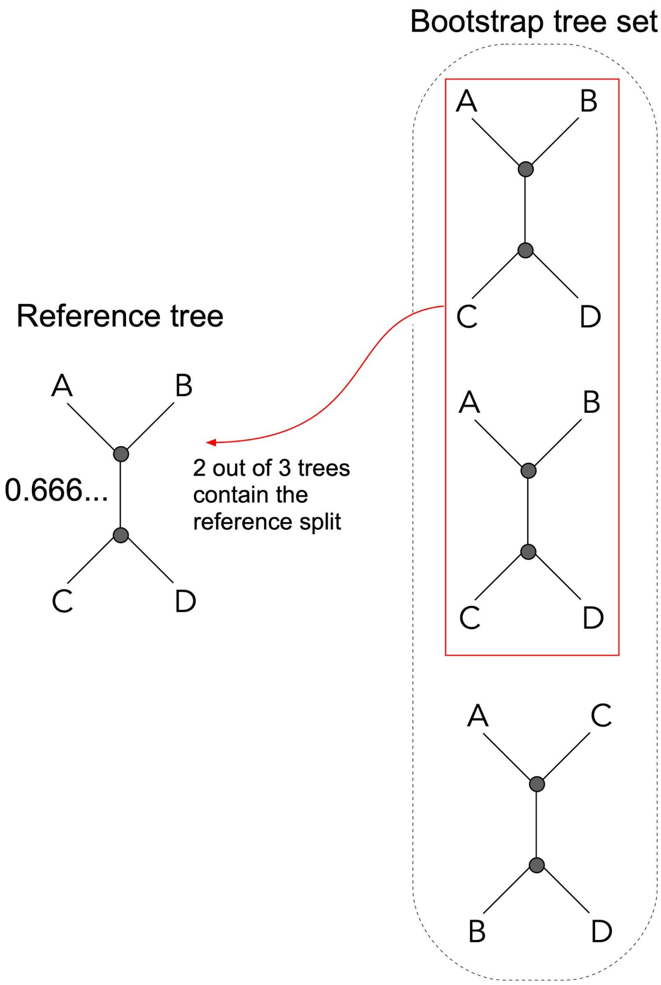


**Figure S16. Phylogenetic bootstrap calculation.** Outline of bootstrap proportion/branch support computations. For each inner branch in the reference tree on the left, we count how frequently the bipartition/split (*AB*|*CD* in the example) occurs in the set of bootstrap trees on the right. Here the bipartition/split/branch is present in two out of three bootstrap trees, and hence the branch support is ⅔.

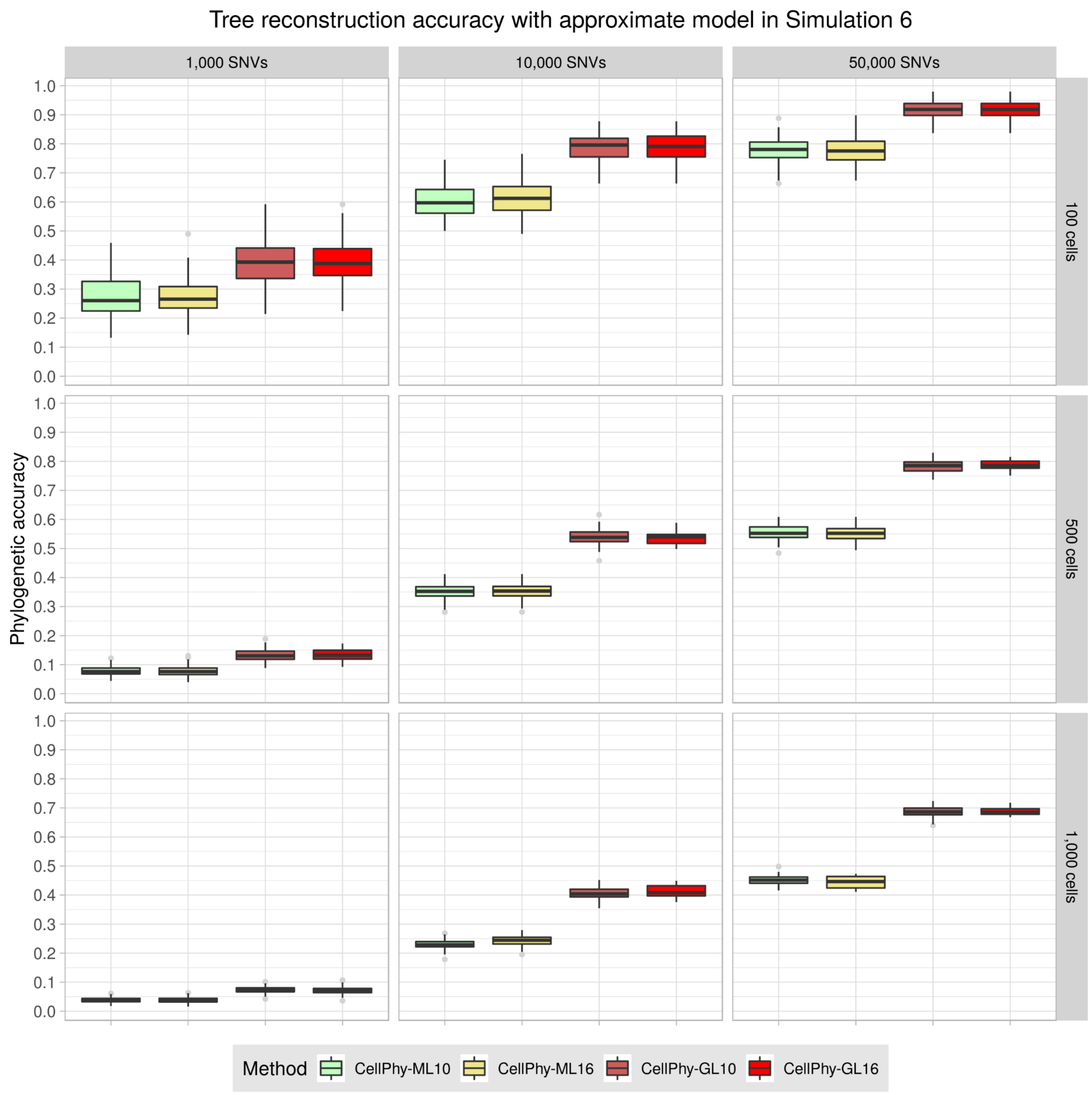


**Figure S17. Phylogenetic accuracy with an approximate model in Simulation 6 ("NGS-large")**. The approximate 10-state model for unphased genotypes (ML10/GL10) yields very similar accuracy to the original 16-state model (ML16/GL16). This observation holds for all simulations in our study (data not shown). Data simulated under mutational signature S1 and with a 5x sequencing depth. Phylogenetic accuracy is defined as 1 – nRF (see Methods). See Figure S1 for an explanation of the boxplots.

### Supplementary Note 1. Genotype error model

P (*N* | *M*) is the probability of observing the single-cell genotype *N* after sequencing, given the true genotype *M*, diploid and biallelic. We consider two types of technical errors that can result in a wrong genotype, allelic dropout (ADO) and amplification/sequencing error (ERR), which occur at rates $\delta$ and $\varepsilon$, respectively. Note that we allow for the presence of both ADO and ERR in the same observed genotype.

Allelic dropout occurs during single-cell whole-genome amplification (scWGA) when one of the two alleles is not amplified and cannot be represented in the observed data. ADO implies a single allele, as otherwise the genotype is "missing". Thus, the rate $\delta$ is the probability that the amplification of one or the other allele has failed, and therefore that we observe the homozygous genotype defined by the amplified allele. Given the phased genotype a|b, and with "_ " indicating the dropped allele:

$$\text{ADO rate } (\delta) = P\ (\_|b \mid a|b) + P\ (a|\_ \mid a|b)$$

An amplification/sequencing error occurs when the observed allele is not the true allele. Given that the ERR rate tends to be small, we assume a maximum of one ERR per genotype. Specifically:

$$\text{ERR rate } (\varepsilon) = P\ (b|a), \text{ where } b \mathrel{!=} a$$

#### Phased genotypes error model

Under these assumptions, there are seven scenarios (I-VII) with non-zero probability for the calculation of P (*N* | *M*) for phased genotypes, with alleles *a*-*d*:

*If the true genotype is homozygous*

I. $P\ (a|a \mid a|a) = (1 - \delta) * (1 - \varepsilon) + \delta * (1 - \tfrac{1}{2}\varepsilon) = 1 - \varepsilon + \tfrac{1}{2}\delta\varepsilon$

For example, P (AA | AA). This can happen in two ways: (1) without ADO (i.e., $1 - \delta$) and without ERR (i.e., $1 - \varepsilon$), or (2) after ADO in either allele ($\delta$) and without ERR in the non-dropped allele (i.e., $1 - \tfrac{1}{2}\varepsilon$).

II. $P\ (a|b \mid aa) = (1 - \delta) * \tfrac{1}{2}\tfrac{1}{3}\varepsilon = (1 - \delta) * \tfrac{1}{6}\varepsilon$

For example, P (AT | AA). This can only happen without ADO (i.e., $1 - \delta$), followed by an ERR in a specific allele, that is converted to one of the other three nucleotides (i.e., $\tfrac{1}{2}\tfrac{1}{3}\varepsilon$).

III. $P\ (b|b \mid a|a) = \delta * \tfrac{1}{2}\tfrac{1}{3}\varepsilon = \tfrac{1}{6}\delta\varepsilon$

For example, P (TT | AA). Because we ignore more than one ERR per genotype, this can only happen through ADO in either allele (i.e., $\delta$), followed by an ERR in the non-dropped allele (i.e., $\tfrac{1}{2}\tfrac{1}{3}\varepsilon$).

For a given true homozygous genotype, we have 1 I, 6 II, 3 III combinations, so the sum of probabilities is: $1 - \varepsilon + \tfrac{1}{2}\delta\varepsilon + \varepsilon - \delta\varepsilon + \tfrac{1}{2}\delta\varepsilon = 1$

*If the true genotype is heterozygous*

IV. P ($a|a$ | $a|b$) = $(1 – δ) * ½ ⅓ ε + ½ δ * (1 – ½ ε) + ½ δ * ½ ⅓ ε = ½ δ + ⅙ ε – ⅓ δε$

For example, P (AA | AT). This can happen in three ways: (1) without ADO (i.e., 1 – δ) and with ERR in allele j (i.e.,½ ⅓ ε), (2) after ADO in allele j (i.e., ½ δ) followed by no ERR in allele i (i.e., 1 – ½ ε), or (3) after ADO in allele i (i.e., ½ δ) followed by an ERR in allele j (i.e., ½ ⅓ ε).

V. P ($c|c$ | $a|b$) = $δ * ½ ⅓ ε = ⅙ δε$

For example, P (CC | AT). Because we ignore the possibility of two ERR in the same genotype, this can only occur if there is ADO in either allele (i.e., δ), followed by an ERR in the non-dropped allele (i.e., ½ ⅓ ε).

VI. P ($a|c$ | $a|b$) = $(1 – δ) * (½ ⅓ ε) = (1 – δ) ⅙ ε$

For example, P (AC | AT). This can only happen without ADO (i.e., 1 – δ), through an ERR in allele j (i.e., ½ ⅓ ε).

VII. P ($a|b$ | $a|b$) = $(1 – δ) * (1 – ε)$

For example, P (AT | AT). This case can only happen without ADO (i.e., 1 – δ) and without ERR (i.e., 1 – ε).

For a given true heterozygous genotype, we have 2 IV, 2 V, 4 VI, and 1 VII combinations, so the sum of probabilities is $δ + ⅓ ε – ⅔ δε + ⅓ δε + (1 – δ) ⅔ ε + (1 – δ) * (1 – ε) = 1$. For the remaining scenarios, given the assumptions of the model, P (*N* | *M*) is zero.

**Unphased genotypes error model**

For unphased genotypes, the error model is identical to the phased case, except for scenario II:

P ($a|b$ | $a|a$) = $(1 – δ) * ½ ⅓ ε = (1 – δ) * ⅓ ε$

For example, P (AT | AA). This can only happen without ADO (i.e., 1 – δ), followed by an ERR in one of the alleles, that is converted to one of the other three nucleotides (i.e., ⅓ ε).

### Supplementary Note 2. Approximate model of evolution for unphased diploid genotypes with ten states

Current techniques for producing scDNA-seq data do not reveal the phase of the genotypes (i.e., we do not know which allele is located in the maternal or paternal chromosome). We have also implemented a specific model for unphased genotypes with only ten states that speed up the calculations. However, for unphased states, the probability of changing between a homozygous and a heterozygous genotype is not reversible, as, in principle, the change from genotype *aa* to genotype *ab* is twice more probable (either allele can change) than from *ab* to *aa* (change can occur only in allele *b*). Considering this asymmetry would result in a non-reversible Q matrix, which would yield the calculation of the tree likelihood much more complex and prohibitively slow, as it would require rooted trees. As a compromised, approximate solution, we implemented a GT10 model based on a reversible Q matrix:

$$
Q_{10} = \begin{array}{c} \\ A/A \\ C/C \\ G/G \\ T/T \\ A/C \\ A/G \\ A/T \\ C/G \\ C/T \\ G/T \end{array}
\begin{array}{c}
\begin{array}{cccccccccc} A/A & C/C & G/G & T/T & A/C & A/G & A/T & C/G & C/T & G/T \end{array} \\
\left(\begin{array}{cccccccccc}
-q_{A/A} & 0 & 0 & 0 & \alpha\pi_{A/C} & \beta\pi_{A/G} & \gamma\pi_{A/T} & 0 & 0 & 0 \\
0 & -q_{C/C} & 0 & 0 & \alpha\pi_{A/C} & 0 & 0 & \kappa\pi_{C/G} & \lambda\pi_{C/T} & 0 \\
0 & 0 & -q_{G/G} & 0 & 0 & \beta\pi_{A/G} & 0 & \kappa\pi_{C/G} & 0 & \mu\pi_{G/T} \\
0 & 0 & 0 & -q_{T/T} & 0 & 0 & \gamma\pi_{A/T} & 0 & \lambda\pi_{C/T} & \mu\pi_{G/T} \\
\alpha\pi_{A/A} & \alpha\pi_{C/C} & 0 & 0 & -q_{A/C} & \kappa\pi_{A/G} & \lambda\pi_{A/T} & \beta\pi_{C/G} & \gamma\pi_{C/T} & 0 \\
\beta\pi_{A/A} & 0 & \beta\pi_{G/G} & 0 & \kappa\pi_{A/C} & -q_{A/G} & \mu\pi_{A/T} & \alpha\pi_{C/G} & 0 & \gamma\pi_{G/T} \\
\gamma\pi_{A/A} & 0 & 0 & \gamma\pi_{T/T} & \lambda\pi_{A/C} & \mu\pi_{A/G} & -q_{A/T} & 0 & \alpha\pi_{C/T} & \beta\pi_{G/T} \\
0 & \kappa\pi_{C/C} & \kappa\pi_{G/G} & 0 & \beta\pi_{A/C} & \alpha\pi_{A/G} & 0 & -q_{C/G} & \mu\pi_{C/T} & \lambda\pi_{G/T} \\
0 & \lambda\pi_{C/C} & 0 & \lambda\pi_{T/T} & \gamma\pi_{A/C} & 0 & \alpha\pi_{A/T} & \mu\pi_{C/G} & -q_{C/T} & \kappa\pi_{G/T} \\
0 & 0 & \mu\pi_{G/G} & \mu\pi_{T/T} & 0 & \gamma\pi_{A/G} & \beta\pi_{A/T} & \lambda\pi_{C/G} & \kappa\pi_{C/T} & -q_{G/T}
\end{array}\right)
\end{array}
$$

In this case, we need to estimate five nucleotide exchangeabilities ($\alpha$ = r(*A*↔*C)*, $\beta$ = r(*A*↔*G), $\gamma$*= r(*A*↔*T), $\kappa$* = r(*C*↔*G), $\lambda$* = r(*C*↔*T); let $\mu$* = r(*G*↔*T) = 1*) and nine stationary unphased genotype frequencies ($\pi_{A/A}$, $\pi_{A/C}$, $\pi_{A/G}$, $\pi_{A/T}$, $\pi_{C/C}$, $\pi_{C/G}$, $\pi_{C/T}$, $\pi_{G/G}$, $\pi_{G/T}$; $\pi_{T/T}$= 1 - $\Sigma\pi_{a/b}$). Regarding the error model, the definitions of ADO, ERR and P (*N* | *M*) are the same as for the GT16, except for P (*a*|*b* | *aa*) = (1 – δ) * ⅓ ε.

In simulations, this (wrong) reversibility assumption did not affect performance; there was no decrease in accuracy, and the calculations were ~2X faster than for the GT16 model (see Figure S17).

### Supplementary Note 3. Standard phylogenetic likelihood calculations on DNA sequence alignments

Let us consider how we compute the phylogenetic likelihood on a standard multiple DNA sequence alignment (MSA). The calculation of the phylogenetic likelihood for our genotype model (see Section 'phylogenetic likelihood' in the main text) is precisely analogous. Given an MSA comprising the sequences under study, a 4x4 instantaneous rate matrix Q, the ability to compute the corresponding 4x4 transition probability matrix $P_t$, and the stationary frequency vector π for the four nucleotides, we calculate the likelihood as follows.

We assume that MSA sites evolve independently of each other. Hence the overall likelihood of the tree, given the MSA, is the product over the per-site likelihoods. Thus, it suffices to consider how to compute the likelihood for a single MSA site. Given a fixed tree topology, with fixed branch lengths and known inner/ancestral states for one MSA site, we simply compute the per-site likelihood as the product over all transition probabilities along the tree branches, times the stationary frequency of the root state. While the ancestral nucleotide states are typically not known, we can still calculate the likelihood of the site as the sum over the per-site likelihoods for all possible assignments of nucleotides (i.e., sum over all possible evolutionary histories that could have generated the data, given the tree) to the ancestral states of the given tree topology.

While this, at first glance, appears to be computationally intense (e.g., for a tree with two ancestral nodes, there already exist 4^2 distinct possible assignments of nucleotides to inner nodes of the tree), the likelihood on such a tree can be efficiently computed via the so-called Felsenstein pruning algorithm. The key idea of this algorithm is to calculate the likelihood bottom-up, that is, from the tips toward the root of the tree, and to store intermediate results, so-called conditional likelihood vectors (CLVs), at each inner node of the tree (Figure S15). CLVs essentially summarize the signal stemming from the subtree they root. That is, they tell us how likely it is to observe an A, C, G, or T, given (conditional on) the subtree they represent. Thus, every inner state in our calculations for one single MSA site consists of a vector containing four conditional likelihoods for, A, C, G, and T, respectively. At the tips, we initialize this vector to (1.0, 0.0, 0.0, 0.0) if we have an A, to (0.0, 1.0, 0.0, 0.0) if we have a C, etc. as the nucleotide state is known and assuming that we are not uncertain about its state.

However, under a single, uniform sequencing error ε, for instance, we can initialize the CLV for A at the tip of the tree as (1.0 - 3ε, ε, ε, ε). This flexibility is used for modeling genotype errors as presented in Section 'single-cell genotype errors' of the main text. The Felsenstein pruning algorithm allows us to compute the likelihood of a given tree, with given branch lengths, and given evolutionary rates specified in Q. Now, to obtain the maximum likelihood (ML) score for such a fixed tree topology, we need to optimize the free parameters of the model, that is, the branch lengths and the rates in Q concerning the likelihood using appropriate numerical optimization routines.

Finally, we also need to find the tree topology with the best ML score, which constitutes an NP-hard optimization problem. In layman's terms, this means that we are simply not able to find the globally best ML tree, as there are too many possible tree topologies (e.g., for 50 sequences, there exist already 283806325080779912837729172696128150920628587998105114415737667754150390625 distinct alternative tree topologies. Furthermore, suppose our model Q is time-reversible. In that case, we can root our tree at any branch or node and will always obtain the identical analytical likelihood score for all possible rooting locations. This is computationally convenient, as, for a given tree, we do not need to determine the optimal root placement to compute its ML score. Hence, the output of phylogenetic inferences under time-reversible models is always an unrooted binary tree, as any rooting will be mathematically meaningless.

Adapting the nucleotide substitution model to a model with more states is, in essence, straight-forward, as precisely the same computational procedure can be used to analyze protein data with 20 states, or our model with ten states here (see corresponding sections on the genotype error and likelihood model in the main text).
